# A Detailed Model for Understanding the Human Neocortex

**DOI:** 10.64898/2026.01.29.702592

**Authors:** Natalí Barros-Zulaica, Lida Kanari, Vishal Sood, Pranav Rai, Alexis Arnaudon, Ying Shi, Darshan Mandge, Werner Van Geit, Michael Zbili, Maria Reva, Elvis Boci, Rodrigo Perin, Maurizio Pezzoli, Ruth Benavides-Piccione, Javier DeFelipe, Eline Mertens, Christiaan de Kock, Idan Segev, Henry Markram, Michael W. Reimann

## Abstract

The neocortex underlies cognitive abilities that set humans apart from other species. Although Ramon y Cajal initiated its study in the 19th century, much about its fundamental properties remain poorly understood. Biologically detail modeling, has been shown to serve as a tool to understand the modeled system better. By comparing computational models for different species we can highlight functional differences between them, find their anatomical or physiological basis and thus improve our understanding of cortical function. In this study we built a detailed model of a human cortical microcircuit following an established workflow. We compared the human data and results against a previously published reconstruction of rat cortical circuitry. To parametrize the human model, we gathered new original data on human morphological reconstructions, axonal bouton densities, single cell and synaptic recordings. We combined them with data available in the literature and open-sourced databases. We also developed various strategies to overcome the missing data, such as generalizing or adapting data from rodents. The resulting model consists of seven columnar units with similar characteristics. Each column has a radius of 476 ***µ***m, a height of 2622 ***µ***m, a volume of 1.86 mm^3^, a total cell density of 24,186 cells/mm^3^, on the order of 35,000 cells, around 12 million connections and approximately 47 million synapses. Comparing the rat and the human model showed that the human cortex is less dense in terms of cell bodies than the rodent cortex. Human cells have more complex branching, but lower bouton densities than rodent cells. However, the number of connections between cell types is similar.

## 1 Introduction

The neocortex is believed to be the brain structure that most strongly differentiates humans from other mammals, as its activity is related to the cognitive skills thought to be uniquely human, such as spoken language, writing, composing music, or inventing complex machines [1, 2].

Since Santiago Ramon y Cajal’s anatomical and histological inquiries in the 19th century [3], many studies have been performed to unravel and understand the mammalian neocortex, most of them in rodents, from all possible perspectives: its anatomical structure [1, 4]; its cell classification from different angles, such as: morphological features [5–7], molecular [8], transcriptomics [9] and single cell physiology [6]; its connectivity [10]; synaptic physiology between pairs of connected cells [10–12] and its network activity [13].

Despite of the many studies about the mammalian neocortex, very little is known about the human neocortex. Due to reasonable ethical constraints, the amount of experimental data in human and primates is limited. Consequently, comparative studies between species are necessary. Many studies have described structural and functional rules that are common between different mammal species, but also found specific characteristic differences between them, e.g., between humans and rodents [1, 4, 10, 11, 14].

Neuroscientific questions that are difficult to address experimentally, have been investigated through computational modeling methods. These methods span a wide range of levels of abstraction, from building simplified and phenomenological model descriptions [15, 16], to morphologically and biophysically detailed computational reconstructions of neuronal circuits [17–20]. By integrating experimental data across multiple scales, computational models enable systematic exploration of circuit mechanisms and generate quantitative, testable predictions that can guide and be validated by future experimental studies.

It has been previously shown that the use of morphologically detailed computational models and simulations can help speed up the process of unraveling the mystery of brain circuits [17, 21, 22]. In this approach, most of the sparse data available within the community is combined and integrated into a coherent model. By comparing computational models for different species we can highlight functional differences between them, find their anatomical or physiological basis and thus improve our understanding of cortical function.

Consequently, in this study we built a biological detailed model of the human temporal cortex following the approach introduced in Markram et al. [17] for the reconstruction of a rat cortical microcircuit. We generated new, original datasets on: morphological reconstructions of excitatory and inhibitory neurons; axonal bouton densities of excitatory cells; and whole cell recordings for excitatory and inhibitory neurons. We integrated these data with existing data from the literature to parameterize the model. To address gaps in the available human data, we also developed and applied several strategies to compensate for missing parameters, including the generalization and adaptation of data from other species.

This study has two primary objectives. First, we aim to systematically summarize the available experimental data on the human cortex and to identify and characterize the remaining gaps in current knowledge. Second, we seek to construct a biologically accurate model of the human cortical microcircuit in order to investigate its structural and functional properties through direct comparison with previously developed rodent models.

## 2 Results

### 2.1 Circuit overview

Most of the human data collected came from temporal cortical areas T20 and T21. We gathered data from *postmortem* and *in vivo* healthy tissue as well as from the literature (see Methods). For those building steps with sufficient human data, the rat data was used to extract biological insights, predictions and conclusions by comparison. However, in the situations were the human data set was insufficient, we used rat data to create new approaches, by finding common rules that would help us extrapolate the missing data from the small data set (Fig.1).

**Figure 1.**
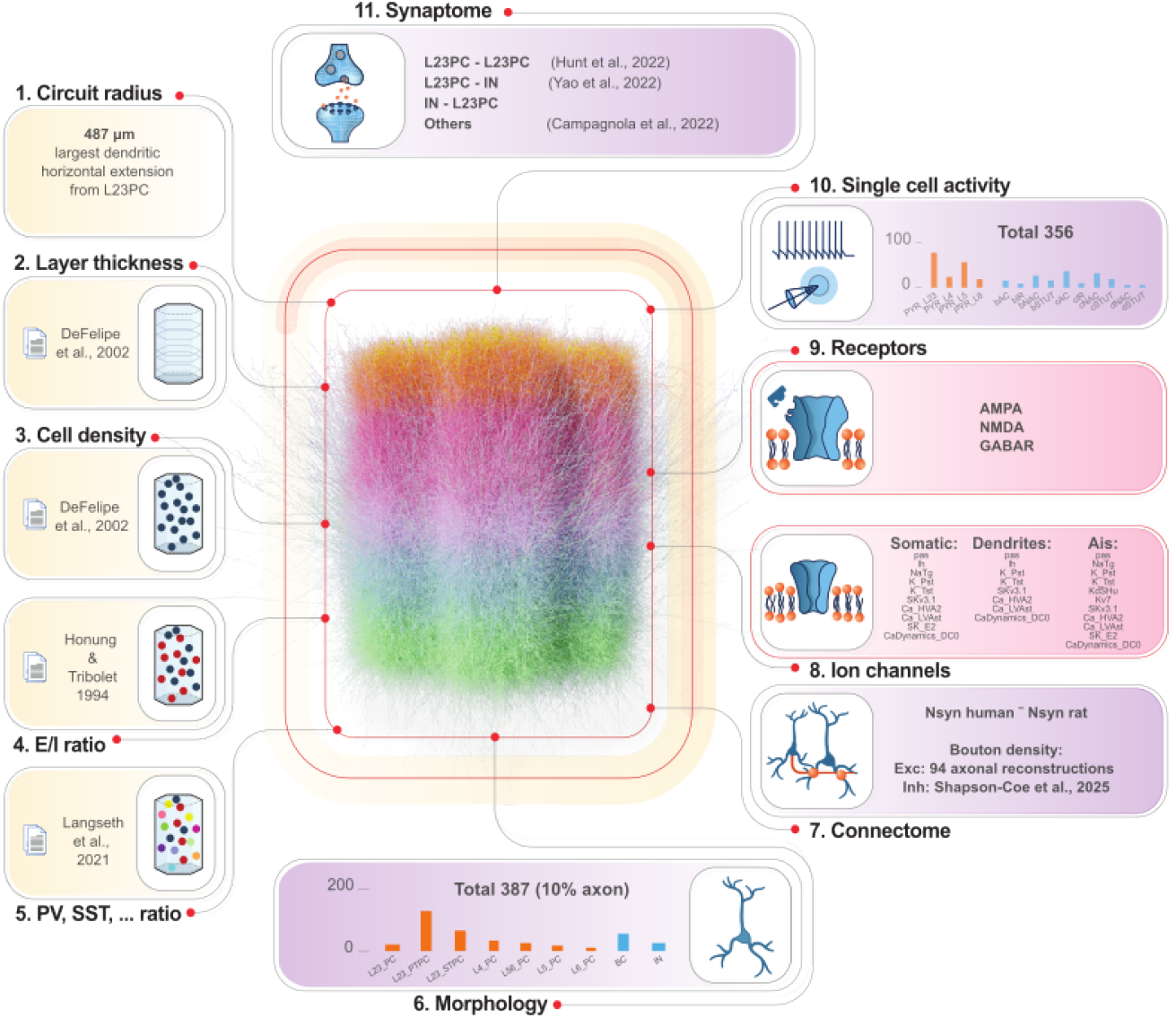
Data collection overview for each step of the model reconstruction pipeline [17]. In the center, visualization of the full circuit with seven columns. Number of cells visualized was reduced to 1% for clarity and neurons were colored by layer. Defining the volume of the circuit required two sets of data circuit radius (1) and cortical layer thicknesses (2). After we have to define how the cells are going to be populating the volume, for that purpose we need to know: cell density (number of cells per mm^3^) per layer (3), ratios of inhibitory vs excitatory cells per layer (4), and the proportion of the different cell types (parvalbumin (PV), somatostatin (SST), …) (5). To populate the circuit we need a collection of morphological reconstructions (6). Then we connect the neurons using data on number of synapses per connection (Nsyn) and axonal bouton densities (7). Now we can implement activity, first for single cells using data on ion channels (8) and receptors (9) and single whole cell recordings (10). And lately recordings between pairs of cells (11).The different colors represent the completeness of data modality: complete (yellow), partially complete (violet), missing (red).

Several of the required datasets for building the circuit were complete and obtained from literature: layer thickness, cell density, ratios of excitatory vs inhibory cells and cell composition (Fig.1, yellow boxes). However, not all required datasets were complete. Data on human ion channels and receptors were substituted completely by rat data (Fig.1, red boxes). Data on morphological reconstructions was partially completed. The majority of the data belonged to layers 2 and 3 excitatory neurons and were didn’t have the axonal branch reconstructed. Consequently, we conclude that more data on morphological reconstructions is required to complete the data set on inhibitory neurons and excitatory neurons from deeper layers, as well as more complete reconstructions including at least a portion of the axon 1, step 6). Connectome data was considered partially complete 1, step 7). Although we found two publications [23, 24] supporting the assumption that the number of synaptic contacts between pairs is similar between rat and human, we consider that more studies would be required in order confirm our assumption. Physiological data on single cells and connected pairs was also considered partially completed 1, steps 10 and 11). Despite of having a decent amount of data, most of the data came from layers 2 and 3, meaning that more data would be necessary for deeper layers.

Despite missing some data, we were able to build a human cortical model with scientific interest. The model is composed of seven columnar units of similar characteristics. Each column was composed by: a radius of 476 *µ*m, a height of 2622 *µ*m, a volume of 1.86 mm^3^, a total cell density of 24,186 cells/mm^3^, on the order of 35,000 cells, around 12 million connections and approximately 47 million synapses (Fig**??**).

### 2.2 Single cell classification and models

#### 2.2.1 Neuronal morphologies

The morphological reconstructions underwent an iterative curation process [17], during which they were corrected and rotated (see Methods). This quality-control step is essential for model building to ensure proper model functionality during simulations. Morphologies were classified based on the different morphological features using a topological classification method [6, 25]. We found five morphological types for excitatory neurons: two types for layer 2 and layer 3 neurons, which were already classified in a previous study [6]. Excitatory cells of deeper layers (layer 4, 5 and 6) were assigned in one cell type per layer due to the sparsity of data (Fig.1, step 6). Subsequently, morphologies were cloned through a cloning process [17] that allowed us to create a collection of morphologies with increased local variability.

Layers could not be accurately assigned to most of the inhibitory reconstructions due the limitations of layer assessment in the human tissue. Inhibitory cells are typically classified based on the shapes of axonal reconstructions [26]. However, due to the lack of sufficiently accurate axonal reconstructions in human inhibitory cells, it was not possible to assign inhibitory cell types accurately. We classified the inhibitory neurons as basquet cells (BC) and other inhibitory neurons (IN).

The topological properties of human and rodent neurons have revealed distinct differences in their branching patterns. Human dendrites of pyramidal cells exhibit a high density of branches around the soma (200 - 500 µm), with branches originating in close proximity to the soma and extending to greater radial distances. This characteristic is consistently observed across cortical layers (2, 3, and 5) [14, 27] and other brain areas such as the hippocampus [28]. These differences are expected to have an impact on the network connectivity [27].

#### 2.2.2 Electrical models

The construction of the single-cell electrical models (e-models) followed a standard two-step process: First, we extracted electric features (e-features) defined in eFEL [29] that encapsulated the relevant information about the shapes of the experimental voltage recordings. Secondly, we performed e-feature optimization to constrain the model’s passive and active parameters, including ion channel conductances and input resistance. The optimization minimizes the error between the experimental (mean) and model e-features values (see Methods).

We classified the neurons into 14 electrical types (e-types (Fig.2) for constructing electrical models (e-models) that were used in the circuit: 4 for pyramidal (PYR, excitatory) neurons with one per each cortical layer (PYR_L23, PYR_L4, PYR_L5, PYR_L6); and 10 for interneurons (IN, inhibitory): (IN_dNAC, IN_cNAC, IN_cAC, IN_bNAC, IN_bAC, IN_bSTUT, IN_cSTUT, IN_dSTUT, IN_cIR and IN_bIR).

**Figure 2.**
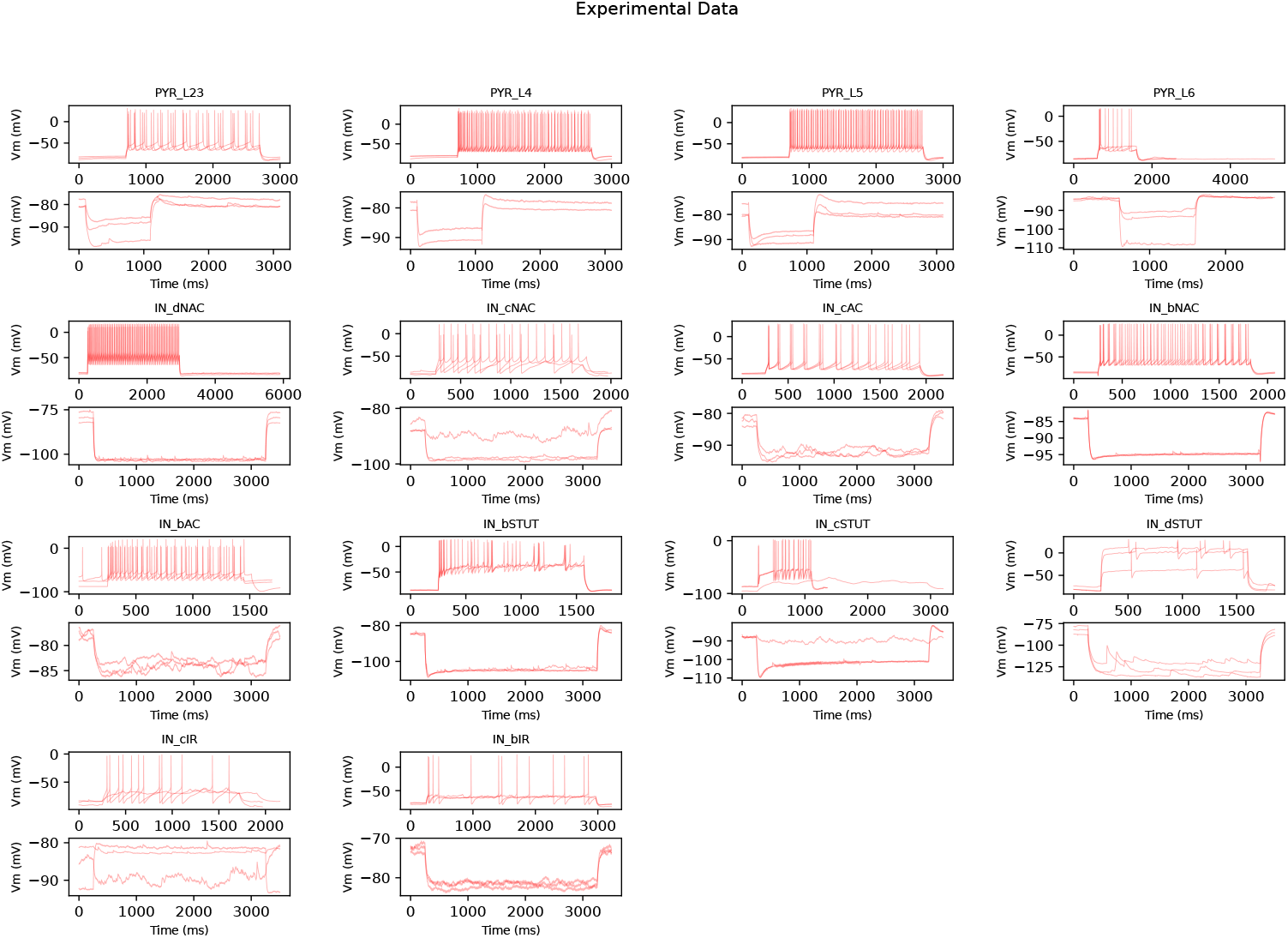
Experimental Data for different electrical types used for e-model construction

We used multi-objective optimization to construct a biophysically realistic e-model for each e-type, using 3D-reconstructed morphologies, electrical features, and ion channel models distributed throughout the neuron. We used the BluePyEModel [30] single-cell modelling workflow to build each e-model. We used CMA-ES (co-variance matrix adaptation evolution strategy)-based multi-objective optimisation implemented in BluePyOpt [31] for model optimisation. E-models were validated with somatic electrophysiological protocols not used for optimization. Ion channel models were distributed in the soma, axon initial segment, and the apical & basal dendrites (Fig.4).

We also compared the e-features extracted from experimental data of human and rat for pyramidal neurons for various layers [32] (Fig.3). The AP amplitude, AP width, input resistance, time constant, and rheobase revealed differences in the active and passive neuronal properties of the two species. The spikecount vs stimulus current plot (Fig.3, bottom) reveals that the spiking behavior also differs. Between human and rat pyramidal neurons.

**Figure 3.**
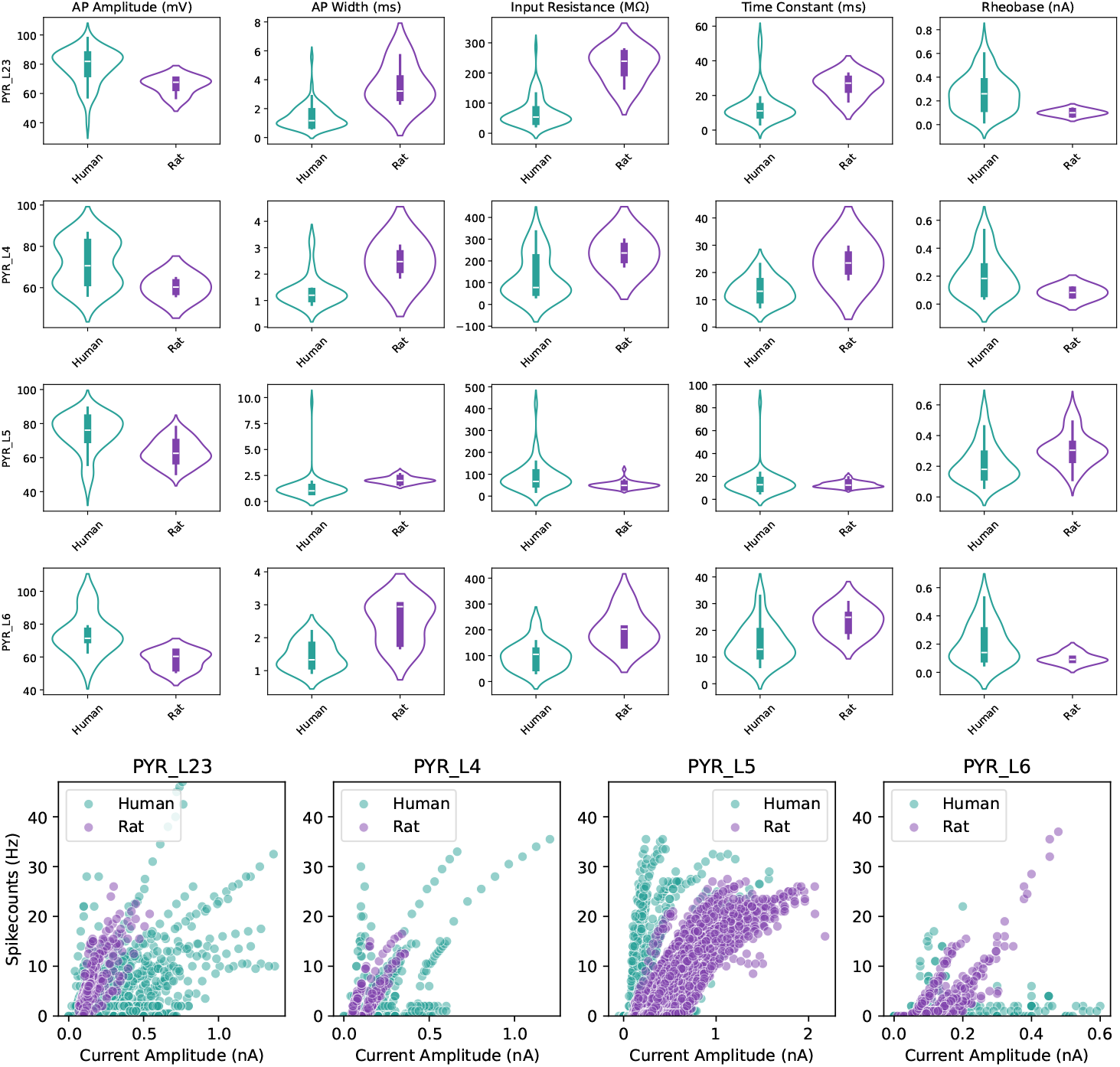
Human and Rat Experimental Data comparison: (Top) The AP amplitude, AP width, input resistance, time constant, and rheobase for various pyramidal neuron electrophysiology data for human (teal) and rat (purple). (Bottom) Spikecount vs. Stimulus Current plot for the pyramidal neuron data for human (teal) and rat (purple). These comparisons reveal very different properties for the two species.

The e-models were in good agreement with the experimental data, as seen in the comparison of experimental and simulated features (Fig.4C1-C4), traces (Fig.**??**), and low z-scores for optimisation features.

**Figure 4.**
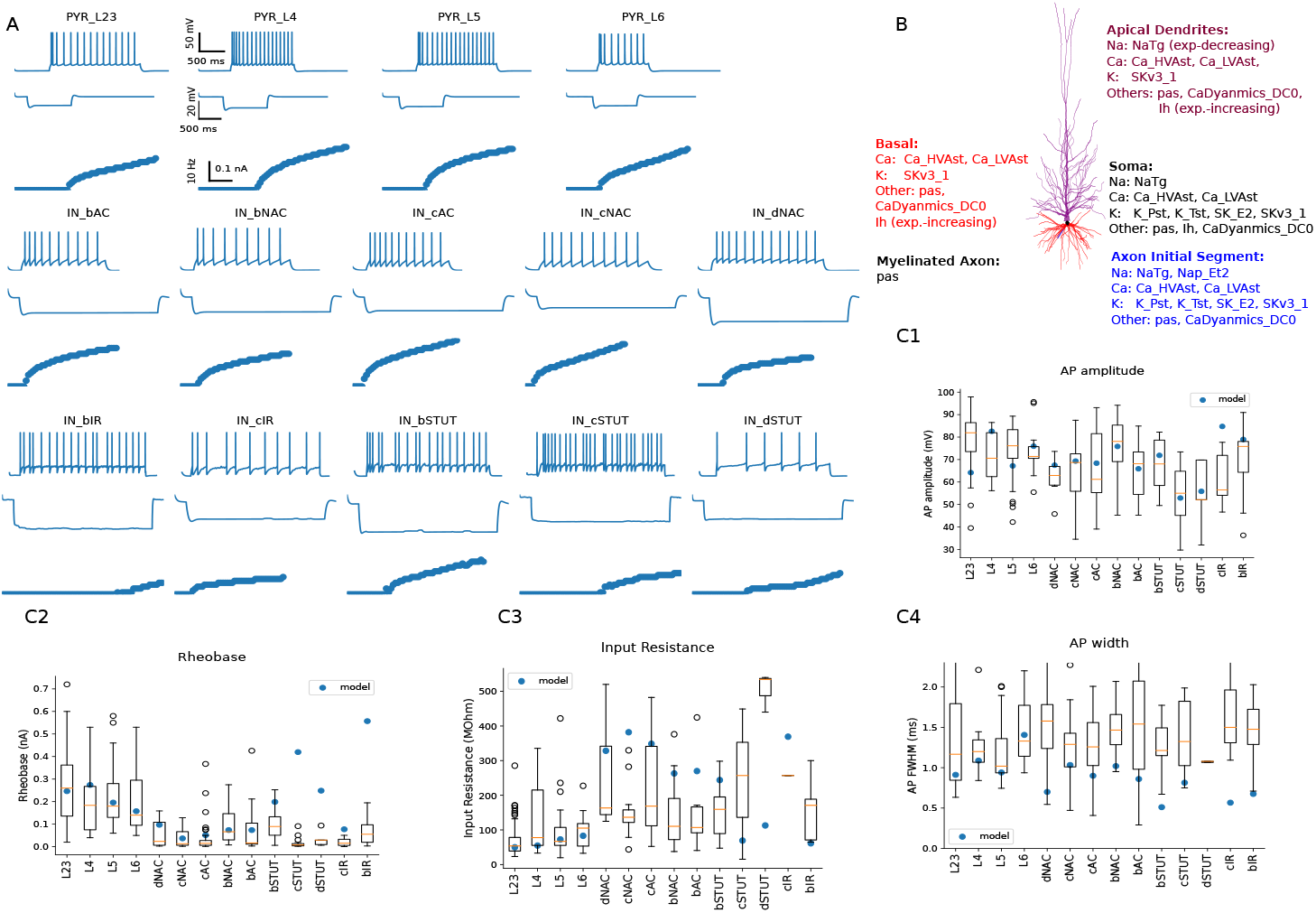
Electrical Models used in the circuit. A. E-model responses for excitatory (PYR_L23, PYR_L4, PYR_L5, PYR_L6) and 10 interneurons (inhibitory) interneuron e-types [5] for : (IN_dNAC, IN_cNAC, IN_cAC, IN_bNAC, IN_bAC, IN_bSTUT, IN_cSTUT, IN_dSTUT, IN_cIR, IN_bIR). Each e-model panel in A shows response for small depolarising (150/140 %) (top), hyperpolarising (IV_- 100, middle), and frequency-current (FI) curve for step current injections. B. A PYR L4 model, morphology showing model ion channel distribution in different sections of the reconstructed neuron morphology. C1-C4 shows the model (blue circle) and experimental feature (box) comparison for AP amplitude, rheobase, input resistance, and AP width. L23, L4, L5, and L6 labels correspond to the respective PYR e-models.

Finally, we generalise the above electrical models to all placed morphologies in the circuit. For this, we used the standard strategy of Markram et al. 2015 [17, 32], which consists of evaluating all possible pairs (also called combos) of cloned morphologies (m-models) and e-models, to keep only the ones for which their scores are acceptable. In Fig.5 we show the results of the model management (see Methods) as the number of pairs of morphologies and models that pass the criteria for use in populating the circuit. We see that for most interneuronal types, all combos pass, except for two STUT models. For pyramidal cells, most models have around 80% pass, except for layer 4. For this model, the rheobase of many cells is large, and the resulting spiking behavior too far from the recordings. This is likely a result of a non-optimal choice of exemplar morphology for optimisation, which could be improved in future versions to improve the morphological variability of this cell type.

**Figure 5.**
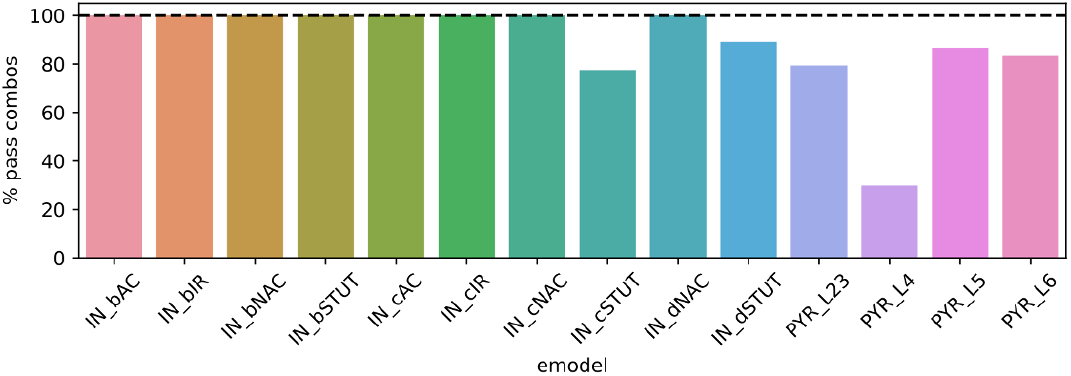
Fraction of pairs of morphologies and models (or combos) which pass the model management selection for the microcircuit.

### 2.3 Circuit anatomical structure

The next steps after having a set of morphological models to populate the circuit are: First, define the dimensions of the circuit. In this occasion we were building a circuit that was composed by seven columns or circuit units in the shape of hexagons, as it was done in our reference publication [17]. For that we needed to define the radius of the circuit and the thickness of the cortical layers. Second, establish cell distributions and proportions for each layer as in cell density, ratios of excitatory and inhibitory neurons, and the proportions of cell subtypes (e.i. parvalbumin, somatostatin, …). Third, once the neurons are placed, we determined their connectivity. For that we used an algorithm [33] (see Methods) that required mostly data on number of synapses between pairs (Nsyn) and axonal bouton density per cell type. We will describe each of them in detail on the following subsections.

#### 2.3.1 Circuit volume: radius and layer thickness

We computed the column radius following the same approach used in Markram et al. [17]. Through this mean we found that the minimal radius where the density of dendrites saturates at the center was 349.40 µm (which was approximately 1.7 times wider than the rat circuit built in 2015). However, as we wanted to ensure the representation of all the morphological reconstructions in our model, we decided to increase the column radius, as few (n=3) of our layer 3a excitatory morphologies had a total horizontal extension of 952 µm (Fig.6A), which lead to a column radius of 476 µm (2.3 times wider than the rat circuit built in 2015).

**Figure 6.**
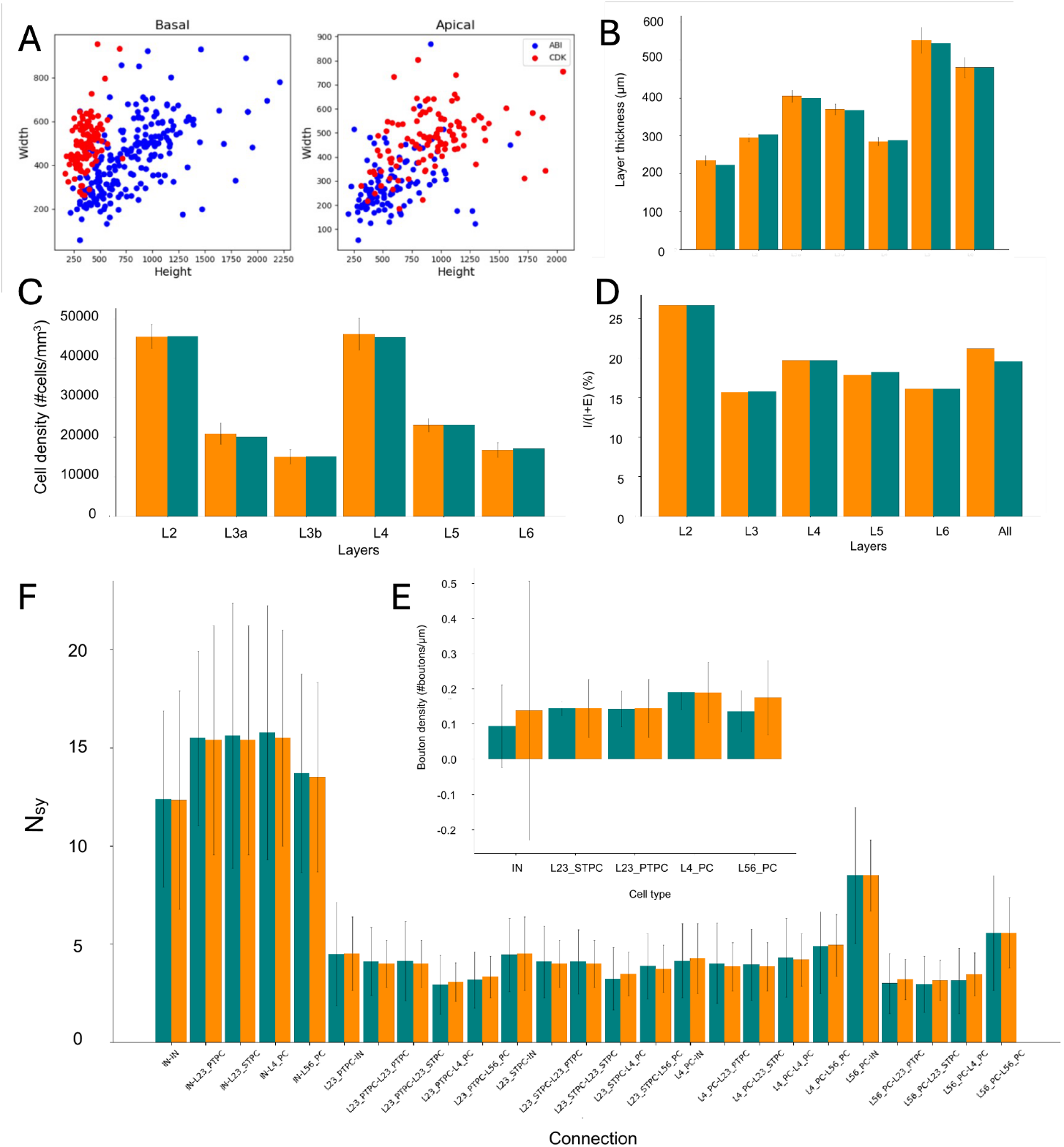
(A) Horizontal dendritic extension for apical and basal dendrites of the set of pyramidal neuronal reconstructions from two different sources. Circuit structural validations for:(B) layer thickness, (C) cell density, (D) ratios of inhibitory vs excitatory cells. Circuit connectivity data validations for: (E) bouton densities per cell type and (G) Number of synapses per pairs.

Regarding layer thickness, we used the data published in DeFelipe et al., 2002 [4] (Fig.6B), which is in consensus with more recent publications [23, 34], and it averages data from three subjects for all cortical layers, while the others studied fewer subjects (n=1, [34]) and did not provide as much granularity for thicker layers, as layer 3 [23].

#### 2.3.2 Cell distribution within the circuit volume

Similarly to the data for the layer thickness, cell density, cell excitatory-inhibitory ratios and cell composition data were collected from the literature. For cell densities (number of neurons per mm^3^), we used the data published in DeFelipe et al. [4] (Fig.6C). These values were corroborated by more recent publications [1, 34].

For cell excitatory-inhibitory ratios per layer we used the data described in table 1 of Hornung and Tribolet 1994 [35], because we considered this study more robust than recent ones, mainly due to the number of subjects (n=4 vs n=1 respectively) (Fig.6D).

**Table 1.** Summary of synaptic parameters across cortical connections.

| Connection | con. prob (%) | Gsyn (nS) | Nsyn | PSP amp (mV) | $\bar{U}$ | $D$ (ms) | $F$ (ms) | Source |
| --- | --- | --- | --- | --- | --- | --- | --- | --- |
| L23_PC - L23_PC | 0.14 | $0.47 \pm 0.1$ | $4 \pm 1.2$ | $1.36 \pm 1.19$ | $0.53 \pm 0.2$ | $319.72 \pm 156.83$ | $38.42 \pm 95.01$ | Hunt et al. [24] |
| L23_PC - L23_IN (SST) | 0.19 | $0.38 \pm 0.1$ | $8 \pm 1$ | $2.16 \pm 1.35$ | $0.09 \pm 0.01$ | $140 \pm 10$ | $670 \pm 10$ | Yao et al. [38] |
| L23_IN - L23_IN | 0.125 | 2.26 | 12.33 | $-1.9 \pm 0.5$ | $0.32 \pm 0.1$ | $689.15 \pm 575.28$ | $3268.85 \pm 3825.73$ | Campagnola et al. [10] |
| L6_PC - L6_PC | 0.09 (L5) | 1.94 | 3.23 | $0.84 \pm 0.1$ | $0.28 \pm 0.1$ | $446.63 \pm 206.03$ | $4339.37 \pm 4885.71$ | Campagnola et al. [10] |
| L6_PC - L6_IN | 0.09 (L5) | 1.0 | 8.5 | $0.86 \pm 0.1$ | $0.14 \pm 0.1$ | $843.88 \pm 582.01$ | $3322.44 \pm 5553.63$ | Campagnola et al. [10] |
| L5_PC - L23_PC | 0.14 | 0.68 | 3.14 | $0.46 \pm 0.1$ | $0.44 \pm 0.1$ | $108.53 \pm 65.26$ | $217.93 \pm 194.70$ | Campagnola et al. [10] |
| L5_PC - L5_PC | 0.07 | 1.94 | 5.57 | $0.89 \pm 0.5$ | $0.43 \pm 0.1$ | $865.77 \pm 2061.05$ | $1660.51 \pm 3237.73$ | Campagnola et al. [10] |
| L5_PC - L5_IN | 0.09 | 1.0 | 8.5 | $2.20 \pm 0.5$ | $0.18 \pm 0.1$ | $632.83 \pm 600.27$ | $5506.56 \pm 6266.20$ | Campagnola et al. [10] |
| L5_IN - L5_PC | 0.09 | 1.0 | 13.5 | $-2.17 \pm 0.5$ | $0.47 \pm 0.1$ | $135.01 \pm 64.37$ | $8.94 \pm 13.04$ | Campagnola et al. [10] |
| L5_IN - L4_PC | 0.36 | 1.95 | 15.5 | $-0.30 \pm 0.1$ | $0.53 \pm 0.1$ | $241.16 \pm 40.53$ | $1300.54 \pm 1838.51$ | Campagnola et al. [10] |
| L4_PC - L4_PC | 0.007 | 0.58 | 4.18 | $1.88 \pm 0.5$ | $0.19 \pm 0.1$ | $2705.96 \pm 4863.57$ | $3363.75 \pm 4588.72$ | Campagnola et al. [10] |
| L4_PC - L4_IN | 0.3 | 0.58 | 3.5 | $2.1 \pm 0.5$ | $0.5 \pm 0.1$ | $275.85 \pm 129.43$ | $62.39 \pm 70.35$ | Campagnola et al. [10] |
| L4_IN - L4_PC | 0.36 | 0.58 | 15.5 | $-0.12 \pm 0.1$ | $0.33 \pm 0.1$ | $1062.48 \pm 999.65$ | $1463.11 \pm 2128.98$ | Campagnola et al. [10] |
| L4_IN - L4_IN | 0.125 | 0.58 | 12.33 | $-1.8 \pm 0.5$ | $0.25 \pm 0.1$ | $1906.43 \pm 2828.03$ | $1648.08 \pm 3660.55$ | Campagnola et al. [10] |
| L23_IN - L23_PC | 0.19 | $1.24 \pm 0.1$ | $12 \pm 1$ | $-1.05 \pm 0.44$ | $0.3 \pm 0.1$ | $1300 \pm 100$ | $2 \pm 1$ | Yao et al. [38] |

#### 2.3.3 Circuit anatomical connectivity

##### Number of synapses between connected cell pairs (Nsyn)

Surprisingly, data in literature, mainly from two publications [24, 36], showed that the number of synapses between cell types in the human cortex is similar than those from rodent (mouse and rat) cortex. From these studies we obtained the following values for the following connections: layer 2/3 pyramidal cell to layer 2/3 inhibitory cell (3.3±1.5 for human; 2.9±1.5 for rat) [36]. Layer 2/3 PC to layer 2/3 PC (4.0±1.2 for human; 3.2±1.3 for mouse) [24].

Considering the large number of missing data for the human cortex on these type of data, and the results from the publications, we decided to fill in the lack of human data with rat data on Nsyn.

##### Axonal bounton density per cell type

Numbers of boutons and axonal lengths were computed from 94 dendro-axonal morphological reconstructions of excitatory cells in all layers, similarly to how it was done in Benavides-Piccione et al. [14] (see Methods) (Fig.6F).

To obtain axonal bouton densities for inhibitory cells, we used the data based described in Shapson-Coe et al. [37] (see Methods). From these dataset we selected randomly 5000 segmentations of axons per layer. From those scheleton files, we were able to extract the distances between boutons and total length of the shaft (Fig.6F).

### 2.4 Structural comparative analysis: human model vs rat model

As it was explained before, learning about the human cortex is challenging due to the lack of data. However, comparing species is a valid approach to obtain rules and patterns that could help us extrapolate the missing data.

Considering this perspective, our main results can be summarized as follows: the human cortex is marginally thicker than the rat cortex (1.26×; 2622 µm vs. 2082 µm), with the largest difference in layer 3 (775 µm vs. 353 µm) [1, 4] . Neuronal density is substantially lower in humans (24,186 neurons/mm3) than in rats (109,217 neurons/mm3), a 0.22× difference [1, 4]. The proportion of inhibitory to excitatory cells per layer is higher in humans (21.1% vs. 13%), reflecting a 1.6× increase in inhibitory cell prevalence [35].

Using these species-specific anatomical data and hundreds of unique reconstructed morphologies (see Table**??** for data sources), we built morphologically detailed circuit models following the process described in Markram et al. [17]. The connectivity of these models was predicted as described in Reimann et al. [33]: as a biologically sustainable subset of the axo-dendritic appositions. We compared the models (human and rat) and found differences in their inter- and intra-laminar connection probabilities (Fig.7C1 (human) and C2 (rat)).

**Figure 7.**
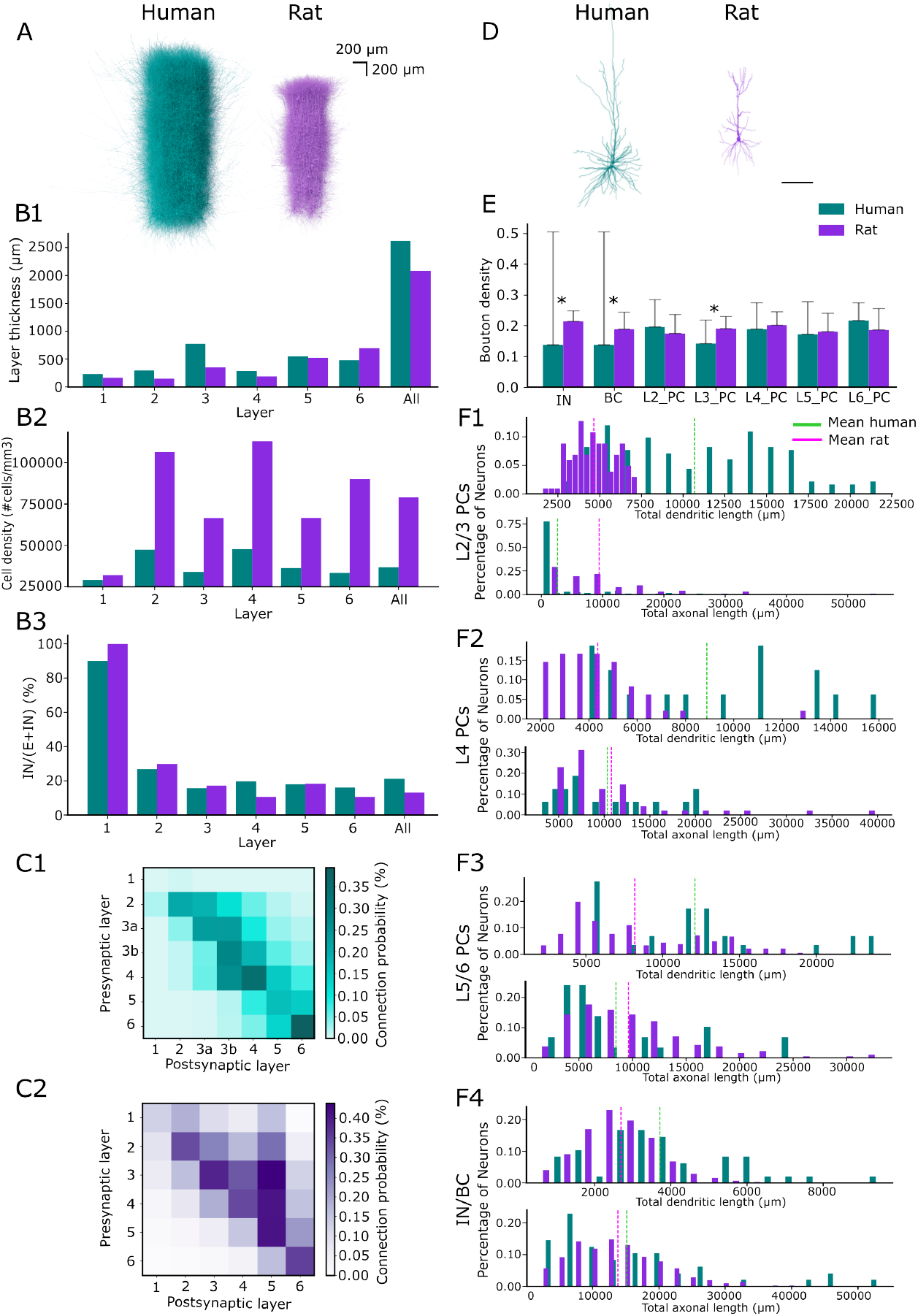
(A) Images of the cortical microcircuit models built for each specie: human (left, teal) and rat (right, purple). (B) Bar plots with values from literature on both species for: layer thickness (B1), cell density (B2) and proportion of inhibitory vs excitatory cells (B3). (C) Connection probability matrices for all neurons separated by layers computed for the computational models: human (top, C1) and rat (bottom, C2). (D) Morphological reconstructions of L3 pyramidal cells in the temporal cortex of the human (left, teal) and the somatosensory cortex of the rat (right, purple).(E) Bar plot showing data on bouton densities for different cell types. (*) points to significant differences when p *<* 0.05. (F) Bar plots showing the percentage of neurons with certain total dendritic (top) and axonal (bottom) lengths for layers 2 and 3 (F1), layer 4 (F2) and layers 5/6 (F3) pyramidal cells and inhibitory and basket cells (F4).

Due to the lack of human data, the human model does not have the same granularity as the rat model concerning morphological cell types. Thus, to compare them we decided to group the cells by layer. Furthermore, most of the human morphological data was gathered for excitatory neurons in layers 2 and layer 3 (≈400), with fewer reconstructions in other layers (≈100) and even fewer for inhibitory neurons (Table**??**). While in the rat the highest connection probabilities were between layers 3, 4 and 5 with layer 5 (0.438, 0.408 and 0.410 respectively), in the case of the human the highest connection probabilities were found in recurrent connections of layers 6 and 4 (0.394, 0.343 respectively; Fig.**??** A and B for values).

In general, the connection probability of the human model was lower than that of the rat. Due to the lack of data on human inhibitory neurons, we also computed the connection probabilities of only the excitatory cells per layer (Fig.**??** (bottom)). In the excitatory subcircuits, we observed that the values of the human connection probability matrix were closer to the ones of the rat in general, while maintaining the qualitative differences outlined above. These differences could be highlighting the fact that in the human temporal cortex few layer 5 pyramidal neurons have an apical dendrite that projects up into layers 2 or 3, while most terminate in layer 4 as suggested by Mohan et al. [23]. In addition, it has been suggested before that remarkable morpho-dendritic differences between human and rodent pyramidal cells (Fig.7D), as seen in Benavides-Piccione et al. [14, 28], could affect the connectivity of the network [27].

We also explored the differences between species at the neuronal level. Data on number of synapses between cell type connections were considered the same between species as no significant differences have been reported in the literature till present [12, 23]. Most local axonal bouton densities were similar between the species, both for inhibitory and excitatory cells. But a significant difference (p *<* 0.05) could be found regarding layer 3 pyramidal cells (human: 0.139 ± 0.077 boutons/*µ*m; rat: 0.187 ± 0.043 boutons/*µ*m; p = 0.000063) and inhibitory cells (human IN and BC: 0.137 ± 0.367; rat IN: 0.198 ± 0.059; rat BC: 0.193 ± 0.063; p(IN) = 0.0098; p(BC) = 0.012) (Fig.7E). It is important to remark that the largest data set of axonal reconstructions in the human belonged to L3 PC (n = 51; against n(L2_PC) = 9, n(L4_PC) = 6, n(L5_PC) = 22 and n(L6_PC) = 2). Thus, more significant differences may emerge as additional data become available.

Regarding morphological reconstructions, we found human dendrites to be approximately twice as large in terms of total length of pyramidal neurons than rat dendrites, but human local axons to be slightly shorter than rat axons. However, it must be noted that these metrics depend on the completeness of reconstructed morphologies, are limited by the technical approach (see methodological considerations). As our human and rat morphological reconstructions came from different experimenters using different instruments and from different labs, their degree of completeness varies.

Overall, the main structural differences showed that neuron density in human cortex is lower with more inhibitory cells than the rodent cortex. However, the number of contact points between pairs of cells remains similar between species, indicating that human cells probably compensate the lack of cell bodies by growing larger and more complex dendritic and axonal branches, as it has been previously studied [27].

### 2.5 Synaptic models

To reconstruct the synaptic physiology we used data gathered through experiments, from literature and from open-sourced data bases.

For connections between layers 2 and 3 pyramidal cells we performed whole-cell patch clamp recordings. We obtained data for 32 connections that were used on a previous study [24]. From this publication we learned that in the human temporal cortex the excitatory postsynaptic potential showed higher amplitude, less failure rate and less variability between protocol repetitions, comparing to mouse temporal cortex. We also observed that the use of synaptic efficacy was higher for human and that the human synapses recovered faster from short-term plasticity protocols than the mouse synapses.

Data for connections between layers 2 and 3 pyramidal-inhibitory and inhibitory-pyramidal connections were obtained from Table S7 of Yao et al. [38].

Data for the rest of the remaining connections were collected from the Allan Brain Institute data base [10]. They provided 1237 synaptic experiments, from 176 different pairs across layer 1 to layer 6.

Synaptic models were built by fitting to the Tsodyks-Markram synaptic models [39–41] the peaks of the postsynaptic potentials from the synaptic recordings (see Methods).

## 3 Discussion

In this study we have reported the available human data required to build a biological detailed model of the human temporal cortex following the approach described by Markram et al. [17] (Fig.1). We saw that anatomical data such as layer thickness, density of neurons per layer, ratios of excitatory and inhibitory neurons per layer, as well as data on cells proportions per layer are available from the literature. Data on structural connectivity for Nsyn showed similar values for both species. However, very little references (only two [24, 36]) reported this assumption. We consider that more data on putative synapses between pairs of neurons would be necessary to corroborate our assumption. Regarding neuronal morphological reconstructions and recordings on single cell and pairs of cells, most of the data came from superficial layers 2 and 3 excitatory neurons, while data on deeper layers (layers 4 to 6) and inhibitory cells were sparser. The vast majority of missing data were for ion channels and receptors. Although there are data available on potassium voltage-gated (Kv) and hyperpolarization-activated cyclic nucleotide-gated channels (HCN) [42], when trying to combine these mechanisms with rat ion channel mechanisms within the same single human neuron, its behavior became very complex and difficult to validate, so we decided to keep all ion channel and receptor mechanisms from rat and we encourage the community to keep working on gathering these sort of important data.

The summarized data in Fig.1 were used to build the human cortical circuit described in this article. This model is an early prototype that would be updated with new data. By building it, we aimed to highlight the missing data sets as well as understand the human cortex better in its different levels. As it has been previously seen, computational models of this sort have been used as a tool to understand different brain areas such as the somatosensory cortex [17, 21, 22], the hippocampus [43] or the thalamus [44]. We can also use these type of models to study different brain abilities as neuromodulation [45] or plasticity [46]. Other studies have followed the approach of building similar models of only layers 2 and 3 of the human temporal cortex to study network connectivity [27], or even depression [38, 47].

By building our model we have learned that bouton densities of layer 3 pyramidal cell show significant difference difference between rat and human (Fig.7E), while no significant differences where observed for other layers. Moreover, data found in the literature describing number of synapses between pyramidal cells, also showed similar values for both species [12]. Additionally, anatomical data obtained through electron-microscopy images showed larger number of synapses per neuron in the human compared to rodent [1], but lower number of synapses per volume unit [4]. We also observed that the human cortex has lower density of cell bodies, higher proportion of inhibitory neurons, compared to the rodent (Fig.7B2 and B3) and that human cortical pyramidal cells have larger (Fig.7F) and more complex dendritic arborization [27] than pyramidal cells from rodents. Intuitively, combining all the previous facts, we could hypothesize that human pyramidal neurons seem to fill the empty space between them with more complex branching, that leads to more synapses per cell, but lower synaptic density (in volume) compared to rodent. This supports the studies of Franz Nissl [48] and Constantin Von Economo [49] that showed higher density of neurons in dog compared to human and a higher capability of interactions between neurons that were widely separated, respectively. That led Cragg [50] to hypothesize that lower soma densities provide the space required for more complex branching, which enrich neuronal interaction, therefore a higher number of synapses per neuron could be expected [1].

Differences in connection probability between layers were also observed. The low values for layer 1 in the human model (Fig.7C1), reflect the great sparsity of human data regarding inhibitory neurons compared to the rat. However, considering only the excitatory cells lead to connectivity matrices with comparable values (Fig **??**). We found big differences between the models: recurrent connections in layers 4 and 6, for which connection probability was higher than that of rat and higher connection probability for rat layer 5 cells connecting recurrently and with layer 4 and 3. This result is probably affected by the lack of human morphological data on deeper layers .

In summary, in this study we pointed out to the main groups of missing data necessary to build a biological detailed model of the human temporal cortex and we concluded with interesting results. Nevertheless, in this study we present a primary version of the model, in the future we would keep exploding the model with simulations and implementing new data sets.

## 4 Methods

### 4.1 Experimental data

#### 4.1.1 Ethical statements

##### CSIC

Human brain tissue was obtained at autopsy from the Unidad Asociada Neuro-max—Laboratorio de Neuroanatomía Humana, Facultad de Medicina, Universidad de Castilla-La Mancha, Albacete, Spain, and Laboratorio Cajal de Circuitos Corticales Universidad Politécnica de Madrid-Consejo Superior de Investigaciones Científicas (CSIC), Madrid, Spain. The tissue was obtained following national laws and international ethical and technical guidelines on using human samples for biomedical research purposes.

##### INF

All procedures were performed with the approval of the Medical Ethical Committee of the Vrije Universiteit Medical Centre (protocol 2012/362) and in accordance with Dutch license procedures and the Declaration of Helsinki. Written informed consent was provided by all subjects for the data and tissue used for scientific research. All data was anonymized.

##### LNMC

All procedures were performed with the approval of the “Commission Cantonale d’éthique de la recherche sur l’être humain” of the Canton of Vaud (protocol 449/14) and in accordance with Swiss license procedures and the Declaration of Helsinki. Written informed consent was provided by all subjects for the data and tissue used for scientific research. All data was anonymized.

#### 4.1.2 Morphological reconstructions

All neurons were subjected to stringent quality control, including evaluation of staining and visualization, as well as inspection for slicing-related artifacts (e.g., sectioning planes misaligned with the apical dendritic axis). Only cells that met these criteria were digitally reconstructed using Neurolucida software (MicroBrightField, Williston, VT, USA) with a 100× oil-immersion objective. Detailed reconstruction procedures are described in Benavides-Piccione et al. [14] and Hunt et al. [24].

### 4.2 Bouton density data

#### 4.2.1 Computed from excitatory neurons

The Neurolucida software was used to extract some morphological variables from some of the reconstructed morphologies. Axonal varicosity density was defined as a a swelling of the axon exceeding the typical variation in diameter of the adjacent axonal shafts per axonal length (see Benavides-Piccione et al. [14] for more details).

#### 4.2.2 Computed from axonal segmentations of inhibitory cells

We estimated bouton densities of inhibitory neurons by analyzing data available in the H01 open-sourced data set [37]. To this end we downloaded 5000 scheleton files of axons, randomly selected, for all layers. Using the software provided (python code and jupyter notebooks), we selected the axonal segments that belonged to inhitory cells and counted the number of root points in it. A root point is the point in the shaft from which the bouton (axonal varicosity) grows. We computed the total length of the axonal segment. Consquently, bouton density was computed as number of root point divided by the total length of the axonal segment.

### 4.3 Electrophysiological Recordings

#### 4.3.1 Slice preparation

##### INF

All experimental procedures were approved by the Medical Ethics Committee of the Vrije Universiteit Medical Center (VUmc) and conducted in accordance with Dutch regulatory requirements and the Declaration of Helsinki. Written informed consent was obtained from all patients. Human cortical tissue was collected from the middle temporal gyrus (MTG) during neurosurgical procedures, where non-pathological tissue was removed to access deeper pathological regions.

Following resection, tissue samples were immediately transferred to a sealed container containing ice-cold artificial cerebrospinal fluid (aCSF). This solution consisted of (in mM): 110 choline chloride, 26 NaHCO_3_, 10 D-glucose, 11.6 sodium ascorbate, 7 MgCl_2_, 3.1 sodium pyruvate, 2.5 KCl, 1.25 NaH_2_PO_4_, and 0.5 CaCl_2_. The aCSF was continuously bubbled with carbogen (95% O_2_ / 5% CO_2_), and the pH was adjusted to 7.3 using 5 M HCl. Transport from the operating room to the neurophysiology laboratory was completed within 20 minutes.

Upon arrival, tissue blocks were rinsed to remove residual blood. After identifying the pia-to–white matter axis, the pia mater was carefully removed. Coronal slices (350 *µ*m thick) were then prepared using a vibratome (Leica V1200S) in ice-cold aCSF. The slices were allowed to recover in a heated bath at 34 °C for 15–30 minutes, followed by an additional recovery period of 1 hour at room temperature.

Recordings were performed in aCSF containing (in mM): 125 NaCl, 3 KCl, 1.2 NaH_2_PO_4_, 1 MgSO_4_, 2 CaCl_2_, 26 NaHCO_3_, and 10 D-glucose (300 mOsm), continuously equilibrated with carbogen gas. Patch pipettes were pulled from borosilicate glass (Harvard Apparatus / Science Products GmbH) and filled with an intracellular solution composed of (in mM): 115 K-gluconate, 10 HEPES, 4 KCl, 4 Mg-ATP, 10 phosphocreatine, 0.3 GTP, 0.2 EGTA, and biocytin (5 mg/ml). The solution was adjusted to pH 7.3 with KOH and an osmolarity of 295 mOsm/kg. During whole-cell recordings, neurons were passively filled with biocytin, enabling subsequent morphological staining and analysis (see methodology in Hunt et al. [24] for more detailed information).

##### LNMC

All experimental procedures were approved by the Swiss Research Ethics Committee and conducted in compliance with Swiss regulatory standards. Written informed consent was obtained from all participants. Human cortical tissue was collected from the idle temporal cortex during neurosurgical interventions, in which non-pathological tissue was removed to access deeper pathological regions.

Immediately after resection, tissue samples were transferred to a sealed container containing ice-cold artificial cerebrospinal fluid (aCSF), which was continuously equilibrated with carbogen gas (95% O_2_ / 5% CO_2_) during transport. The time elapsed between surgical removal and arrival at the neurophysiology laboratory did not exceed 60 minutes. Upon receipt in the laboratory, tissue blocks were rinsed to remove residual blood. After identifying the pia–white matter axis, the pia mater was carefully dissected away. Acute slices (300 *µ*m thick) were then prepared using a vibratome (Leica V1200S) in ice-cold aCSF. Slices were allowed to recover in a heated bath at 34 °C for 15–30 minutes, followed by an additional 1-hour recovery period at room temperature.

Electrophysiological recordings were performed in aCSF containing (in mM): 125 NaCl, 2.5 KCl, 1.25 NaH_2_PO_4_, 1 MgCl_2_ · 6H_2_O, 2 CaCl_2_ · 2H_2_O, 25 NaHCO_3_, and 25 D-glucose, with continuous carbogen bubbling. Patch electrodes were pulled from borosilicate glass (Hilgenberg) and filled with an intracellular solution composed of (in mM): 110 K-gluconate, 10 HEPES, 10 KCl, 4 Mg-ATP, 10 Na-phosphocreatine, 0.3 GTP, and biocytin (3 mg/ml). The intracellular solution was adjusted to pH 7.2– 7.3 with KOH and an osmolarity of 270–300 mOsm.

During whole-cell patch-clamp recordings, neurons were passively filled with biocytin, enabling subsequent histological processing and morphological visualization.

#### 4.3.2 e-types classification

Recordings were performed in excitatory and inhibitory neurons of the human temporal cortex. We classified them into different electrical types (e-types). These include the pyramidal neuron e-types for layers: PYR L23, PYR L4, PYR L5 & PYR L6, and interneurons: burst-accommodating (bAC), continuous-accommodating (cAC), burst non-accommodating (bNAC), continuous non-accommodating (cNAC), delayed non-accommodating (dNAC), burst irregular firing (bIR), continuous irregular firing (cIR), burst stuttering bSTUT), continuous stuttering (cSTUT)& delayed stuttering (dSTUT). The following were the counts per e-type for the 356 electrophysiological recordings: PYR L23 (78 recordings), PYR L4 (24), PYR L5 (57), PYR L6 (19), and interneurons: bAC (16), bIR (9), bNAC (27), bSTUT (16), cAC (37), cIR (10), cNAC(32), cSTUT(19), dNAC(6), dSTUT(6). These recordings consist of responses for various depolarising, hyperpolarising, and combined protocols obtained from somatic current injection. These protocols were grouped into different types based on shape, duration, and frequency: IDRest, IV, APThreshold, sAHP, IDhyperpol, APWaveform, Step, sAHP, FirePattern, and Poscheops. Refer to [32] for protocol details. The protocol amplitudes were expressed as a percentage of the rheobase; e.g., IDRest 130 refers to 130% of the rheobase. See [32] for more details. We extracted electrical features (e-features) defined in the eFEL features extraction library [29] from data collected under various electrophysiological protocols (Table2).

### 4.4 Single Neuron Electrical Models

We constructed 14 e-models: 4 pyramidal (excitatory) neurons with one for each of cortical layers 2/3, 4, 5 & 6 (PYR_L23, PYR_L4, PYR_L5, PYR L6) and 10 interneurons (inhibitory) interneuron e-types [5] for : (IN_dNAC, IN_cNAC, IN_cAC, IN_bNAC, IN_bAC, IN_bSTUT, IN_cSTUT, IN_dSTUT, IN_cIR and IN_bIR).

Some fixed models parameters are: temperature = 34 °C, somatic specific membrane capacitance (cm) = 1*µ*F/cm^2^, dendritic cm = 2*µ*F/cm^2^, myelinated axon cm = 0.02*µ*F/cm^2^, axial resistance (Ra) = 100 Ωcm, sodium Nernst potential (ENa) = 50 mV, potassium Nernst potential (Ek) = -90 mV.

We used the following ion channels as described for rat somatosensory cortical neuron models [32]:

- Sodium (Na) Channels: Transient Na (NaTg), Persistent Na (Nap Et2)
- Potassium (K) Channels: Transient (K_Tst), Persistent (K Pst), Kv3.1 (SKv3 1), Delayed-potassium (KdShu2007), Stochastic (StochKv3), Small-conductance Ca-activated (SKCa, SK E2)
- Calcium Channels: High-voltage-activated Ca(Ca_HVA2) and Low-voltage-activated Ca (Ca LVAst)
- Hyperpolarization-activated current (Ih).

The calcium dynamics (CaDyanmics_DC0) mechanism accounts for calcium buffering and calcium removal through the membrane. Ion channels were distributed over different sections of the morphology as follows: Pyramidal neurons (PYR_L23, PYR_L4, PYR_L5 and PYR_L6):

- Apical Dendrites: Passive (pas), CaDyanmics_DC0, Ca_HVAst, Ca_LVAst, NaTg, SKv3_1, Ih (Exponential-increasing distribution)
- Basal Dendrites: pas, CaDyanmics DC0, Ca_HVAst, Ca_LVAst, Ih (Exponential-increasing distribution)
- Soma: pas, CaDyanmics DC0, Ca_HVAst, Ca_LVAst, SKv3_1, SK_E2, K_Pst, K_Tst, Ih, NaTg (exponential-decreasing distribution)
- Axon Initial Segment: pas, CaDyanmics DC0, Ca HVAst, Ca_LVAst, SKv3_1, SK_E2, K_Pst, K_Tst, NaTg, Nap_Et2
- Myelinated Axon: pas

Interneurons: Regular Firing (IN_bAC, IN_bNAC, IN_cAC and IN_cNAC) Interneurons

- Basal Dendrites: Passive (pas), CaDyanmics DC0, Ca HVAst, Ca_LVAst, Ih (exponential-increasing distribution)
- Soma: pas, CaDyanmics DC0, Ca_HVAst, Ca_LVAst, SKv3_1, SK_E2, K_Pst, K_Tst, NaTg, Ih
- Axon Initial Segment: pas, CaDyanmics DC0, Ca_HVAst, Ca_LVAst, SKv3_1, SK_E2, K_Pst, K_Tst, Nap Et2, NaTg
- Myelinated Axon: pas

Delayed non-accommodating type (IN_dNAC) has all channels in regular-firing interneurons, along with KdShu2007 in soma, axon, and basal dendrites. Irregular and Stuttering firing types (IN_bIR, IN_cIR, IN_bSTUT, and IN_cSTUT) have all channels in regular-firing interneurons, along with StochKv3 in the soma, axon, and basal dendrites. Delayed-stuttering type neuron (IN_dSTUT) has all channels in irregular-firing interneurons, as well as KdShu2007 in the soma, axon, and basal dendrites. All channels were uniformly distributed except the ones mentioned above. Channels were corrected for liquid junction potential whenever possible.

The electrical features (e-features) from the eFEL features extraction library [29] used to optimise the model are shown in table 2.

**Table 2.**
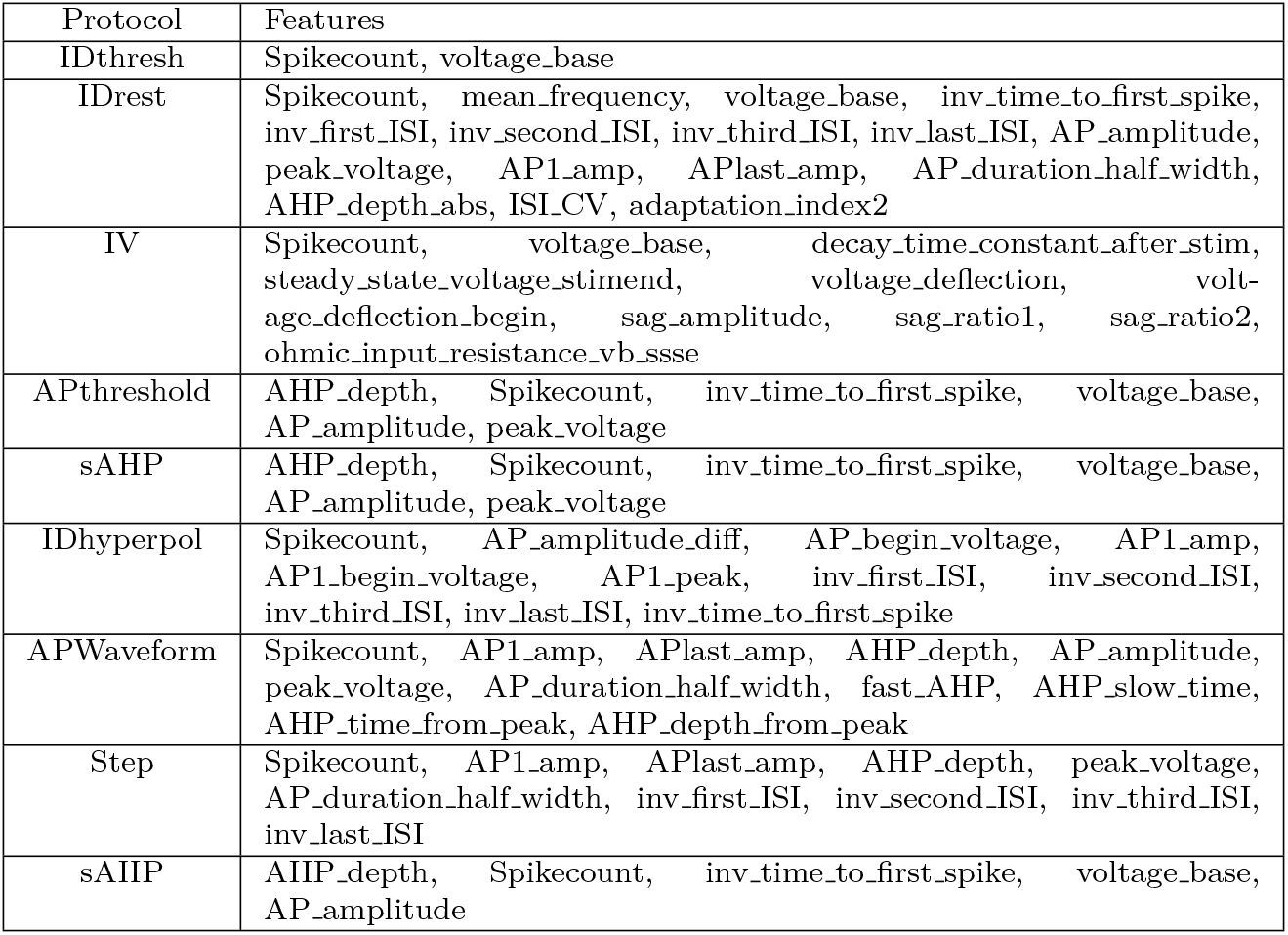
Pyramidal e-model protocol and extracted e-features.

**Table 3.** Interneuron e-model protocols and extracted e-features.

| Protocol | Features |
| --- | --- |
| IDthresh | Spikecount, voltage_base |
| IDrest | Spikecount, mean_frequency, strict_burst_number, strict_burst_mean_freq, voltage_base, inv_time_to_first_spike, inv_first_ISI, inv_second_ISI, inv_third_ISI, inv_last_ISI, AP_amplitude, peak_voltage, APlast_amp, AP_begin_voltage, ISI_CV, ISI_log_slope, adaptation_index2, irregularity_index, number_initial_spikes |
| IV | Spikecount, voltage_base, decay_time_constant_after_stim, steady_state_voltage_stimend, voltage_deflection, voltage_deflection_begin, sag_amplitude, sag_ratio1, sag_ratio2, ohmic_input_resistance_vb_ssse |
| APWaveform | Spikecount, AP1_amp, AP2_amp, AHP_depth, AP_height, AHP_slow_time, AHP_time_from_peak, AP_amplitude, AP_duration_half_width, AP_begin_width |
| sAHP | AHP_depth, Spikecount, inv_time_to_first_spike, voltage_base, AP_amplitude, AP_height |
| FirePattern | strict_burst_number, mean_frequency, AHP_depth, AP_amplitude, AP_height |
| PosCheops | Spikecount |

#### 4.4.1 Optimisation Protocols

We used the following depolarising and hyperpolarising protocols for optimising the models: i) Pyramidal Neurons: IDthresh (for rheobase computation), IDrest: IDrest 150, IDrest_250, where 150 and 250 % of rheobase current, IV: IV_-100, IV_-20, IV_0, APWaveform_300, APthreshold_200, sAHP_200, IDhyperpol_150 and Step: Step_200, Step_300 protocols. ii) Interneurons: IDrest_140, IDrest_250, IV_-100, IV_-40, IV_0, APWaveform_300, sAHP_200, FirePattern_200 and PosCheops_300.

#### 4.4.2 Validation

We validated each e-model after optimisation using protocols that were not used for optimisation. For the pyramidal neuron e-models, we used a protocol to inject multiple random step current injections into the dendrites (apical and basal) and record the firing frequency at the soma. We used a mean somatic frequency e-feature of 10 Hz and a standard deviation of 2 Hz to get an average somatic spike generation from random dendritic stimulation. We use a dendritic stimulus injection of 150 % of somatic rheobase step currents of variable duration and start times. We also validated pyramidal with the following protocols: IDhyperpol_150, sAHP_200, and APthreshold_200. Similarly, we used the randomstepsdendritic_150, sAHP_200, FirePattern_200, and PosCheops_300 models to validate the interneuron models. We used various experimental features from these protocols (as shown in the table) to validate the model by verifying each feature score ¡ or close to 5 experimental standard deviations.

#### 4.4.3 Optimisation Procedure

We used the BluePyEModel [30] single-cell modelling workflow, which reads and extracts electrical features from electrophysiology data, then optimises and validates them to produce the final model. It combines the software eFEL [29], BluePyEfe [51], and Bluepyopt [31]. We use the rheobase-dependent multiobjective optimisation where the protocol amplitudes are scaled based on % of rheobase current. The electrophysiological features extracted from simulated traces are compared with those from experiments to calculate a feature score as follows:

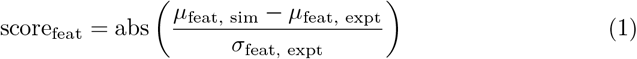

where score_feat_ is a feature score, *µ*_feat, sim_ and *µ*_feat, expt_ are simulation and experimental feature mean and *σ*_feat, expt_ is experimental feature standard deviation. (Refer to [32] for more details.) We replace the reconstructed axon with a stub axon with 60 *µ*m of the axon initial segment and 1000 *µ*m of a myelinated axon. We used the SO-CMA (single-objective covariance matrix adaptation) evolutionary algorithm from BluePyOpt for optimisation for 400 generations, with an offspring size of 20. Some e-models were optimised for fewer optimisation generations, such as when the optimisation stopping criterion is met, as described in BluePyOpt, or for irregular-firing interneurons using stochastic channels, which require comparatively longer optimisation times.

#### 4.4.4 Model Management

We followed and adapted the method described in Markram et al. [17] and Reimann et al. [52] to reconstruct a microcircuit of the human temporal cortex (Fig.1). In brief, we collected 387 morphological reconstructions, belonging to different morphological types. Initial reconstructions were curated to produce a library of Hodgkin-Huxley multicompartimental neuron models. We placed neuron models into the defined volume according to available data on cell densities and composition. We derived the intrinsic connectivity following the method of [33], and assigned synaptic parameters as described in [41]

### 4.5 Connectome

For the digital reconstruction of the connectivity we used the same approach described in Markram et al. [17], Reimann et al. [33]. The required parameters are:

- Neuron density per cortical layer per each morphological type: gathered from literature [4]
- Total axonal length per cell type (key to determine the number of appositions and synapses): obtained from morphological reconstructions
- Mean number of synapses (Nsyn) (mean ± std), per each connected cell pair.
- Connection probability of each type of connection
- Axonal bouton density (mean ± std) per cell type

### 4.6 Synaptic models

The list of necessary parameters per synaptic connection:

- Synaptic conductance (gsyn): normally estimated by fitting post-synaptic (PSP) amplitudes expeirmental values
- Synaptic conductance for the NMDA component (gsynSRSF): multiplier to get the gsyn of the NMDA component
- Number of vesicles ready to be released (NRRP): set 1 in the current model [40]
- coefficient of variation (CV): computed from PSP experimental amplitudes.
- Time decay of AMPA synaptic conductance (dtc): fitting the decay curve to an exponential (in ms)
- Release probability (usyn, U): fitting the experimental trace to the Tsodyks-Markram model [39]
- Time constant of recovery from depression (D): fitting the experimental trace to the Tsodyks-Markram model [39] (in ms)
- Time constant of recovery from facilitation (F): fitting the experimental trace to the Tsodyks-Markram model [39] (in ms)
- Hill curve parameter (uHillCoefficient): to compute it we need PSP data for [Ca2+] between 1 to 4 mM (Fig.S11 from Markram et al. [17]
- Synaptic type: E1 (excitatory, facilitating), E2 (excitatory, depressing), E3 (excitatory, pseudolinear), I1 (inhibitory, facilitating), I2 (inhibitory, depressing) or I3 (inhibitory, pseudolinear) [17] After gsyn calibration, synaptic models validation was performed through *in silico* patch-clamp experiments (Fig.**??**).

## References

[1] DeFelipe, J.: The evolution of the brain, the human nature of cortical circuits, and intellectual creativity. Front Neuroanat (2011) 10.3389/fnana.2011.00029

[2] Jawabri, K.H., Sharma, S.: Physiology, Cerebral Cortex Functions. StatPearls, Treasure Island (FL) (2023)

[3] S.R., C.: Cajal on the Cerebral Cortex. New York: Oxford University Press., ??? (1989)

[4] DeFelipe, J., Alonso-Nanclares, L., Arellano, J.I.: Microstructure of the neocortex: comparative aspects. J Neurocytol. 31, 299–316 (2002)

[5] Ascoli, A.G., Alonso-Nanclares, L., Anderson, S.A., Barrionuevo, G., Benavides-Piccione, R., Burkhalter, A., Buzsáki, G., Cauli, B., DeFelipe, J., Fairén, A., Feldmeyer, D., Fishell, G., Fregnac, Y., Freund, T.F., Gardner, D., Gardner, E.P., Goldberg, J.H., Helmstaedter, M., Hestrin, S., Karube, F., Kisvárday, Z.F., Lambolez, B., Lewis, D.A., Marin, O., Markram, H., Muñoz, A., Packer, A., Petersen, C.C.H., Rockland, K.S., Rossier, J., Rudy, B., Somogyi, P., Staiger, J.F., Tamas, G., Thomson, A.M., Toledo-Rodriguez, M., Wang, Y., West, D.C., Yuste, R., (PING), T.P.I.N.G.: Petilla terminology: nomenclature of features of gabaergic interneurons of the cerebral cortex. Nature Reviews Neuroscience 9(7), 557–568 (2008) 10.1038/nrn2402

[6] Deitcher, Y., Eyal, G., Kanari, L., Verhoog, M.B., Kahou, G.A.A., Mansvelder, H.D., C.P.J. Kock, Segev, I.: Comprehensive morpho-electrotonic analysis shows 2 distinct classes of l2 and l3 pyramidal neurons in human temporal cortex. Cerebral Cortex 27(11), 5398–5414 (2017) 10.1093/cercor/bhx226 https://academic.oup.com/cercor/article-pdf/27/11/5398/33122192/bhx226.pdf

[7] Kanari, L., Ramaswamy, S., Shi, Y., Morand, S., Meystre, J., Perin, R., Abdellah, M., Wang, Y., Hess, K., Markram, H.: Objective morphological classification of neocortical pyramidal cells. Cerebral Cortex (New York, NY) 29, 1719–1735 (2019)

[8] Zeng, H., Sanes, J.R.: Neuronal cell-type classification: challenges, opportunities and the path forward. Nature Reviews Neuroscience 18(9), 530–546 (2017) 10.1038/nrn.2017.85

[9] Yuste, R., Hawrylycz, M., Aalling, N., Aguilar-Valles, A., Arendt, D., Armañanzas, R., Ascoli, G.A., Bielza, C., Bokharaie, V., Bergmann, T.B., Bystron, I., Capogna, M., Chang, Y., Clemens, A., Kock, C.P.J., DeFelipe, J., Santos, S.E.D., Dunville, K., Feldmeyer, D., Fiáth, R., Fishell, G.J., Foggetti, A., Gao, X., Ghaderi, P., Goriounova, N.A., Güntürkün, O., Hagihara, K., Hall, V.J., Helmstaedter, M., Herculano-Houzel, S., Hilscher, M.M., Hirase, H., Hjerling-Leffler, J., Hodge, R., Huang, J., Huda, R., Khodosevich, K., Kiehn, O., Koch, H., Kuebler, E.S., Kühnemund, M., Larrañaga, P., Lelieveldt, B., Louth, E.L., Lui, J.H., Mansvelder, H.D., Marin, O., Martinez-Trujillo, J., Chameh, H.M., Mohapatra, A.N., Munguba, H., Nedergaard, M., Němec, P., Ofer, N., Pfisterer, U.G., Pontes, S., Redmond, W., Rossier, J., Sanes, J.R., Scheuermann, R.H., Serrano-Saiz, E., Staiger, J.F., Somogyi, P., Tamás, G., Tolias, A.S., Tosches, M.A., García, M.T., Wozny, C., Wuttke, T.V., Liu, Y., Yuan, J., Zeng, H., Lein, E.: A community-based transcriptomics classification and nomenclature of neocortical cell types. Nature Neuroscience 23(12), 1456–1468 (2020) 10.1038/s41593-020-0685-8

[10] Campagnola, L., Seeman, S.C., Chartrand, T., Kim, L., Hoggarth, A., Gamlin, C., Ito, S., Trinh, J., Davoudian, P., Radaelli, C., Kim, M.H., Hage, T., Braun, T., Alfiler, L., Andrade, J., Bohn, P., Dalley, R., Henry, A., Kebede, S., Alice, M., Sandman, D., Williams, G., Larsen, R., Teeter, C., Daigle, T.L., Berry, K., Dotson, N., Enstrom, R., Gorham, M., Hupp, M., Dingman, L.S., Ngo, K., Nicovich, P.R., Potekhina, L., Ransford, S., Gary, A., Goldy, J., McMillen, D., Pham, T., Tieu, M., Siverts, L., Walker, M., Farrell, C., Schroedter, M., Slaughterbeck, C., Cobb, C., Ellenbogen, R., Gwinn, R.P., Keene, C.D., Ko, A.L., Ojemann, J.G., Silbergeld, D.L., Carey, D., Casper, T., Crichton, K., Clark, M., Dee, N., Ellingwood, L., Gloe, J., Kroll, M., Sulc, J., Tung, H., Wadhwani, K., Brouner, K., Egdorf, T., Maxwell, M., McGraw, M., Pom, C.A., Ruiz, A., Bomben, J., Feng, D., Hejazinia, N., Shi, S., Szafer, A., Wakeman, W., Phillips, J., Bernard, A., Esposito, L., D’Orazi, F.D., Sunkin, S., Smith, K., Tasic, B., Arkhipov, A., Sorensen, S., Lein, E., Koch, C., Murphy, G., Zeng, H., Jarsky, T.: Local connectivity and synaptic dynamics in mouse and human neocortex. Science, 375–6585 (2022) 10.1126/science.abj5861

[11] Seeman, S.C., Campagnola, L., Davoudian, P.A., al: Sparse recurrent excitatory connectivity in the microcircuit of the adult mouse and human cortex. Elife 7, 37349 (2018)

[12] Hunt, S., Leibner, Y., Mertens, E.J., Barros-Zulaica, N., Kanari, L., Heistek, T.S., Karnani, M.W., Aardse, R., Wilbers, R., Heyer, D.B., Goriounova, N.A., Verhoog, M.B., Testa-Silva, G., Obermayer, J., Versluis, T., Benavides-Piccione, R., Witt-Hamer, P., Idema, S., Noske, D.P., Baayen, J.C., Lein, E.S., DeFelipe, J., Markram, H., Mansvelder, H.D., Schürmann, F., Segev, I., Kock, C.P.J.: Strong and reliable synaptic communication between pyramidal neurons in adult human cerebral cortex. Cereb Cortex 6, 2857–2878 (2023)

[13] Luhmann, H.J.: Dynamics of neocortical networks: connectivity beyond the canonical microcircuit. Pflügers Archiv - European Journal of Physiology 475, 1027–1033 (2023) 10.1007/s00424-023-02830-y

[14] Benavides-Piccione, R., Blazquez-Llorca, L., Kastanauskaite, A., Fernaud-Espinosa, I., Tapia-González, S., DeFelipe, J.: Key morphological features of human pyramidal neurons. Cerebral Cortex (2024) 10.1093/cercor/bhae180

[15] Zhang, L., Zhang, D., Deng, Y., Ding, X., Wang, Y., Tang, Y., Sun, B.: A simplified computational memory model from information processing. Scientific Reports 6, 37470 (2016) 10.1038/srep37470

[16] Wang, T., Wang, Y., Shen, J., Wang, L., Cao, L.: Predicting spike features of hodgkin-huxley-type neurons with simple artificial neural network. Frontiers in Computational Neuroscience (2022) 10.3389/fncom.2021.800875

[17] Markram, H., Muller, E., Ramaswamy, S., al., M.W.R.: Reconstruction and simulation of neocortical microcircuitry. Cell 163, 456–492 (2015)

[18] Billeh, Y.N., Cai, B. L.G.; S., Dai, K., Iyer, R., Gouwens, N.W., Abbasi-Asl, R., Jia, X., Siegle, J.H., Olsen, S.R., Koch, C., Mihalas, S., Arkhipov, A.: Systematic integration of structural and functional data into multi-scale models of mouse primary visual cortex. Neuron (2020) 10.1016/j.neuron.2020.01.040

[19] Gidon, A., Zolnik, T.A., Fidzinski, P., Bolduan, F., Papoutsi, A., Poirazi, P., Holtkamp, M., Vida, I., Larkum, M.E.: Dendritic action potentials and computation in human layer 2/3 cortical neurons. Science 367, 83–87 (2020) 10.1126/science.aax6239

[20] Yao, H.K., Guet-McCreight, A., Mazza, F., Moradi Chameh, H., Prevot, T.D., Griffiths, J.D., Tripathy, S.J., Valiante, T.A., Sibille, E., Hay, E.: Reduced inhibition in depression impairs stimulus processing in human cortical microcircuits. Cell Reports 38(2), 110232 (2022) 10.1016/j.celrep.2021.110232

[21] Isbister, J.B., Ecker, A., Pokorny, C., Bolaños-Puchet, S., Egas Santander, D., Arnaudon, A., Awile, O., Barros-Zulaica, N., Alonso, J.B., Boci, E., Chindemi, G., Courcol, J.-D., Damart, T., Delemontex, T., Dietz, A., Ficarelli, G., Gevaert, M., Herttuainen, J., Ivaska, G., Ji, W., Keller, D., King, J., Kumbhar, P., Lapere, S., Litvak, P., Mandge, D., Muller, E.B., Pereira, F., Planas, J., Ranjan, R., Reva, M., Romani, A., Rössert, C., Schürmann, F., Sood, V., Teska, A., Tuncel, A., Geit, W.V., Wolf, M., Markram, H., Ramaswamy, S., Reimann, M.W.: Modeling and simulation of neocortical micro- and mesocircuitry. part ii: Physiology and experimentation. eLife (2024)

[22] Reimann, M.W., Santander, D.E., Kanari, L., Barros-Zulaica, N.: Neuron morphological physicality and variability define the non-random structure of connectivity. bioRxiv (2025) 10.1101/2025.08.21.671478 https://www.biorxiv.org/content/early/2025/08/22/2025.08.21.671478.full.pdf

[23] Mohan, H., Verhoog, M.B., Doreswamy, K.K., Eyal, G., Aardse, R., Lodder, B.N., Goriounova, N.A., Asamoah, B., Brakspear, A.B.C.B., Groot, C., Sluis, S., Testa-Silva, G., Obermayer, J., Boudewijns, Z.S.R.M., Narayanan, R.T., Baayen, J.C., Segev, I., Mansvelder, H.D., Kock, C.P.J.: Dendritic and axonal architecture of individual pyramidal neurons across layers of adult human neocortex. Cerebral Cortex (New York, NY) 25, 4839–4853 (2015)

[24] Hunt, S., Leibner, Y., Mertens, E.J., Barros-Zulaica, N., Kanari, L., Heistek Karnani, M.M., Aardse, R., Wilbers, R., Heyer, D.B., Goriounova, N.A., Verhoog, M.B., Testa-Silva, G., Obermayer, J., Versluis, T., Benavides-Piccione, R., Witt-Hamer, P., Idema, S., Noske, D.P., Baayen, J.C., Lein, E.S., DeFelipe, J., Markram, H., Mansvelder, H.D., Schürmann, F., Segev, I., Kock, C.P.J.: Strong and reliable synaptic communication between pyramidal neurons in adult human cerebral cortex. Cereb Cortex., 2857–2878 (2023) 10.1093/cercor/bhac246

[25] Kanari, L., D-lotko, P., Scolamiero, M., Levi, R., Shillcock, J., Hess, K., Markram, H.: A topological representation of branching neuronal morphologies. Neuroinformatics 16(1), 3–13 (2018)

[26] Mihaljević, B., Benavides-Piccione, R., Bielza, C., Larrañaga, P., DeFelipe, J.: Classification of gabaergic interneurons by leading neuroscientists. Sci Data 6 (2019) 10.1038/s41597-019-0246-8

[27] Kanari, L., Shi, Y., Arnaudon, A., Barros-Zulaica, N., Benavides-Piccione, R., Coggan, J.S., DeFelipe, J., Hess, K., Mansvelder, H.D., Mertens, E.J., Meystre, J., Campos Perin, R., Pezzoli, M., Daniel, R.T., Stoop, R., Segev, I., Markram, H., de Kock, C.P.J.: Of mice and men: Dendritic architecture differentiates human from mouse neuronal networks. iScience 28(7), 112928 (2025) 10.1016/j.isci.2025.112928

[28] Benavides-Piccione, R., Regalado-Reyes, M., Fernaud-Espinosa, I., Kastanauskaite, A., Tapia-González, S., León-Espinosa, G., Rojo, C., Insausti, R., Segev, I., DeFelipe, J.: Differential structure of hippocampal ca1 pyramidal neurons in the human and mouse. Cerebral Cortex (2020)

[29] Ranjan, R., Geit, W.V., Moor, R., Rössert, C., Riquelme, J.L., Damart, T., Jaquier, A., Tuncel, A., Mandge, D., Kilic, I.: eFEL. 10.5281/zenodo.14222078 . https://doi.org/10.5281/zenodo.14222078

[30] Mandge, D., Jaquier, A., Tanguy, D., Kilic, I., Tuncel, A., Arnaudon, A., Dorp, S.V., Ficarelli, G., Kanari, L., Markram, H., Geit, W.V.: BluePyEModel. https://doi.org/10.5281/zenodo.8283490. 10.5281/zenodo.8283490

[31] Geit, W.V., Gevaert, M., Chindemi, G., Rössert, C., Courcol, J.D., Muller, E.B., Schürmann, F., Segev, I., Markram, H.: Bluepyopt: Leveraging open source software and cloud infrastructure to optimise model parameters in neuroscience. Frontiers in Neuroinformatics (2016) 10.3389/fninf.2016.00017

[32] Reva, M., Rössert, C., Arnaudon, A., Damart, T., Mandge, D., Tuncel, A., Ramaswamy, S., Markram, H., Van Geit, W.: A universal workflow for creation, validation, and generalization of detailed neuronal models. Patterns, 100855 (2023) 10.1016/j.patter.2023.100855

[33] Reimann, M.W., King, J.G., Muller, E.B., Ramaswamy, S., Henry, M.: An algorithm to predict the connectome of neural microcircuits. Frontiers in Computational Neuroscience 9, 1662–5188 (2015)

[34] Shapson-Coe, A., Januszewski, M., Berger, D.R., Pope, A., Wu, Y., Blakely, T., Schalek, R.L., Li, P., Wang, S., Maitin-Shepard, J., Karlupia, N., Dorkenwald, S., Sjostedt, E., Leavitt, L., Lee, D., Bailey, L., Fitzmaurice, A., Kar, R., Field, B., Wu, H., Wagner-Carena, J., Aley, D., Lau, J., Lin, Z., Wei, D., Pfister, H., Peleg, A., Jain, V., Lichtman, J.W.: A connectomic study of a petascale fragment of human cerebral cortex. bioRxiv (2021) 10.1101/2021.05.29.446289

[35] Hornung, J.P., Tribolet, N.D.: Distribution of gaba-containing neurons in human frontal cortex: a quantitative immunocytochemical study. Anat Embryol (Berl). 189, 139–45 (1994)

[36] Molnár, G., Rózsa, M., Baka, J., Holderith, N., Barzó, P., Nusser, Z., Tamás, G.: Human pyramidal to interneuron synapses are mediated by multi-vesicular release and multiple docked vesicles. Elife, 5–18167 (2016)

[37] Shapson-Coe, A., Januszewski, M., Berger, D.R., Pope, A., Wu, Y., Blakely, T., Schalek, R.L., Li, P.H., Wang, S., Maitin-Shepard, J., Karlupia, N., Dorkenwald, S., Sjostedt, E., Leavitt, L., Lee, D., Bailey, L., Fitzmaurice, A., Kar, R., Field, B., Wu, H., Wagner-Carena, J., Aley, D., Lau, J., Lin, Z., Wei, D., Pfister, H., Peleg, A., Jain, V., Lichtman, J.W.: A petavoxel fragment of human cerebral cortex reconstructed at nanoscale resolution. Science (2024) 10.1126/science.adk4858

[38] Yao, H.K., Guet-McCreight, A., Mazza, F., Moradi, H., Prevot, T.D., Griffiths, J.D., Tripathy, S.J., Valiante, T.A., Sibille, E., Hay, E.: Reduced inhibition in depression impairs stimulus processing in human cortical microcircuits. Cell Rep., 110232 (2022) 10.1016/j.celrep.2021.110232

[39] Fuhrmann, G., Segev, I., Markram, H., Tsodyks, M.: Coding of temporal information by activity-dependent synapses. Journal of Neurophysiology 87(1), 140–148 (2002) 10.1152/jn.00258.2001

[40] Barros-Zulaica, N., Rahmon, J., Chindemi, G., Perin, R., Markram, H., Muller, E., Ramaswamy, S.: Estimating the readily-releasable vesicle pool size at synaptic connections in the neocortex. Front Synaptic Neurosci. (2019) 10.3389/fnsyn.2019.00029

[41] Ecker, A., Romani, A., Sáray, S., Káli, S., Migliore, M., Falck, J., Lange, S., Mercer, A., Thomson, A.M., Muller, E., Reimann, M.W., Ramaswamy, S.: Data-driven integration of hippocampal ca1 synaptic physiology in silico. Hippocampus 30, 1129–1145 (2020)

[42] Ranjan, R., Khazen, G., Gambazzi, L., Ramaswamy, S., Hill, S.L., F, F.S., Markram, H.: Channelpedia: an integrative and interactive database for ion channels. Front Neuroinform 30(5), 36 (2011) 10.3389/fninf.2011.00036

[43] Romani, A., Antonietti, A., Bella, D., Budd, J., Giacalone, E., Kurban, K., al.: Community-based reconstruction and simulation of a full-scale model of the rat hippocampus ca1 region. PLoS Biol 22(11) (2024) 10.1371/journal.pbio.3002861

[44] Litvak, P., Hartley, N.D., Kast, R., Feng, G., Fu, Z., Arnaudon, A., Hill, S.L.: Biophysical modeling of thalamic reticular nucleus subpopulations and their differential contribution to spindle dynamics. iScience 28(9) (2025) 10.1016/j.isci.2025.113393

[45] Colangelo, C., Shichkova, P., Keller, D., Markram, H., Ramaswamy, S.: Cellular, synaptic and network effects of acetylcholine in the neocortex. Front. Neural Circuits 13 (2019) 10.3389/fncir.2019.00024

[46] Chindemi, G., Abdellah, M., Amsalem, O., Benavides-Piccione, R., Delattre, V., Doron, M., Ecker, A., Jaquier, A.T., King, J., Kumbhar, P., Monney, C., Perin, R., Rössert, C., Tuncel, A.M., Geit, W.V., DeFelipe, J., Graupner, M., Segev, I., Markram, H., Muller, E.B.: A calcium-based plasticity model for predicting long-term potentiation and depression in the neocortex. Nature Communications 13(1), 3038 (2022) 10.1038/s41467-022-30214-w

[47] Yao, H.K., Mazza, F., Prevot, T.D., Sibille, E., Hay, E.: Spine loss in depression impairs dendritic signal integration in human cortical microcircuit models. iScience 28 (2025) 10.1016/j.isci.2025.113996

[48] Nissl, F.: Nervenzellen und graue substanz. Munch. med. Wschr. 45, 988– 9921023102910601062 (1898)

[49] Vértes, P.E., Alexander-Bloch, A.F., Gogtay, N., Giedd, J.N., Rapoport, J.L., Bullmore, E.T.: Ein koeffizient fur die organisationshohe der grosshirnrinde. Klin. Wsch., 593–595 (1926)

[50] Cragg, B.J.: The density of synapses and neurones in the motor and visual areas of the cerebral cortex. Journal of Anatomy 101(4), 639–54 (1967)

[51] Rössert, C., Geit, W.V., Iavarone, E., Bologna, L.L., Damart, T., Jaquier, A., Mandge, D., Tuncel, A., Kilic, I., Sanin, A., Davison, A.: BluePyEfe. https://doi.org/10.5281/zenodo.8283490. 10.5281/zenodo.12800737

[52] Reimann, M.W., Bolaños-Puchet, S., Courcol, J.-D., Egas Santander, D., Arnaudon, A., Coste, B., Delemontex, T., Devresse, A., Dictus, H., Dietz, A., Ecker, A., Favreau, C., Ficarelli, G., Gevaert, M., Hernando, J.B., Herttuainen, J., Isbister, J.B., Kanari, L., Keller, D., King, J., Kumbhar, P., Lapere, S., Lazovskis, J., Lu, H., Ninin, N., Pereira, F., Planas, J., Pokorny, C., Riquelme, J.L., Romani, A., Shi, Y., Smith, J.P., Sood, V., Srivastava, M., Geit, W.V., Vanherpe, L., Wolf, M., Levi, R., Hess, K., Sch”urmann, F., Muller, E.B., Ramaswamy, S., Markram, H.: Modeling and simulation of rat non-barrel somatosensory cortex. part i: Modeling anatomy. bioRxiv (2022) 10.1101/2022.08.11.503144 https://www.biorxiv.org/content/early/2022/08/15/2022.08.11.503144.full.pdf

