## Supplemental information for "A Detailed Model for Understanding the Human Neocortex"

### 1 Supplementary Information

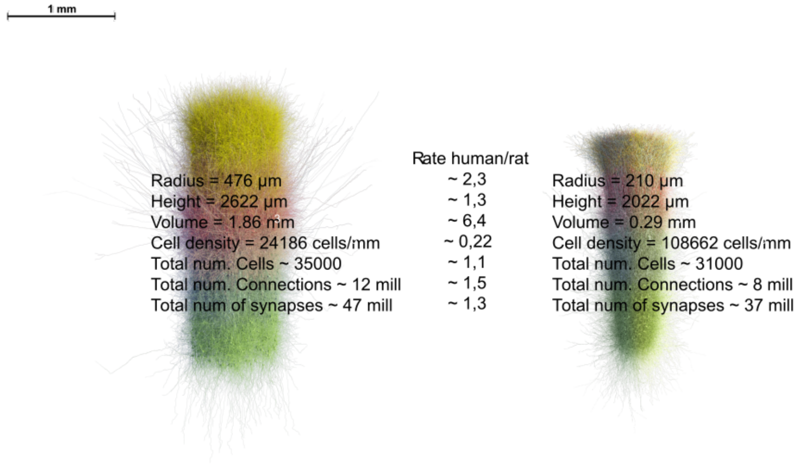


**Figure S1** Comparison between columnar units of human (left) and rat (right) cortical models

**Table S1** References to the human data obtained from literature and their cortical area.

| **Data modality** | **Reference** | **Cortical area** |
|---|---|---|
| Layer thickness ($\mu m$) | DeFelipe 2002 [? ] | Temporal T20, T21 |
| Cell density (num.cells/$mm^3$) | DeFelipe 2002 [? ] | Temporal T20, T21 |
| IN/(EXC+IN) (%) | Honung and Tribolet 1994 [? ] | Prefrontal and Temporal |
| Bouton density (PCs) num. boutons/$\mu m$ | Benavides-Piccione et al., 2024 [? ] | Temporal T20, T21 |
| Bouton density (INs) num. boutons/$\mu m$ | Shapson-Coe et al., 2024 [? ] | Temporal |

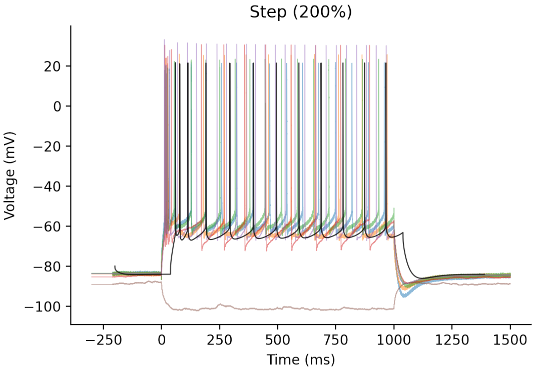


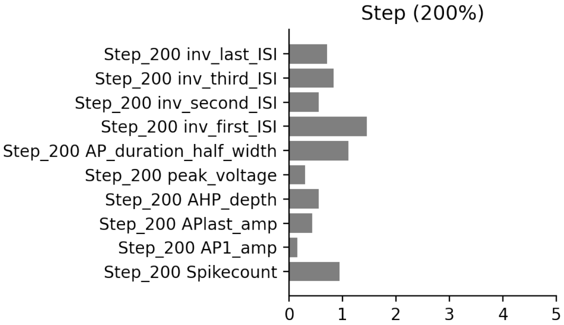


**Figure S2** E-Model vs Experimental Data comparison: (Top) Model trace (black) for Step_200% protocol with experimental (coloured) traces showing their similarity to each other. (Bottom) E-feature score comparison for the optimised e-model features. Lower values indicate that the model feature value is close to the mean experimental feature value.

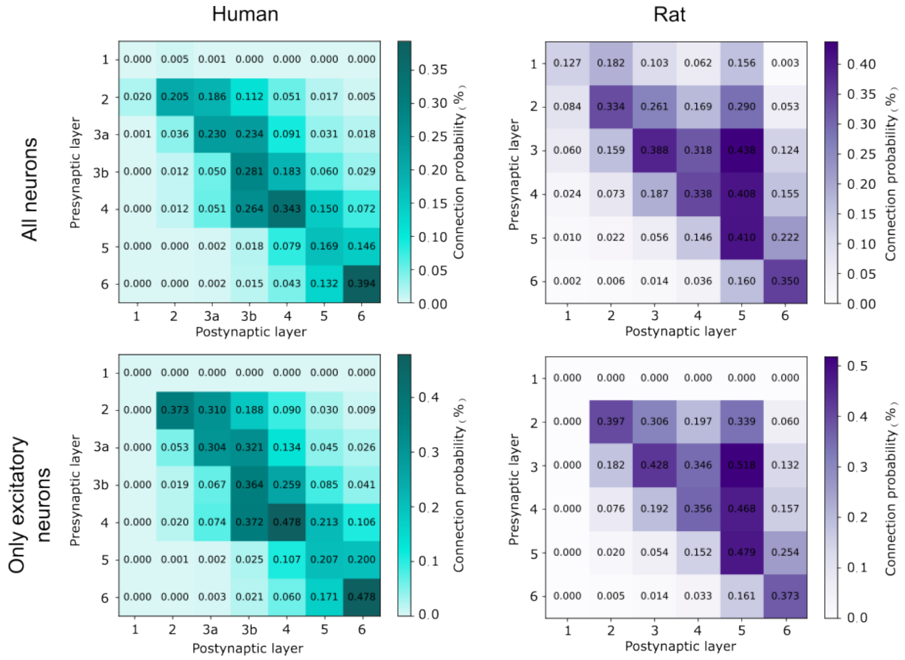


**Figure S3** Connectivity matrices with values for the connection probability, organized by layer for Human (teal) and Rat (purple) for all neurons ( 5. A and B) and for only excitatory cells (C and D)

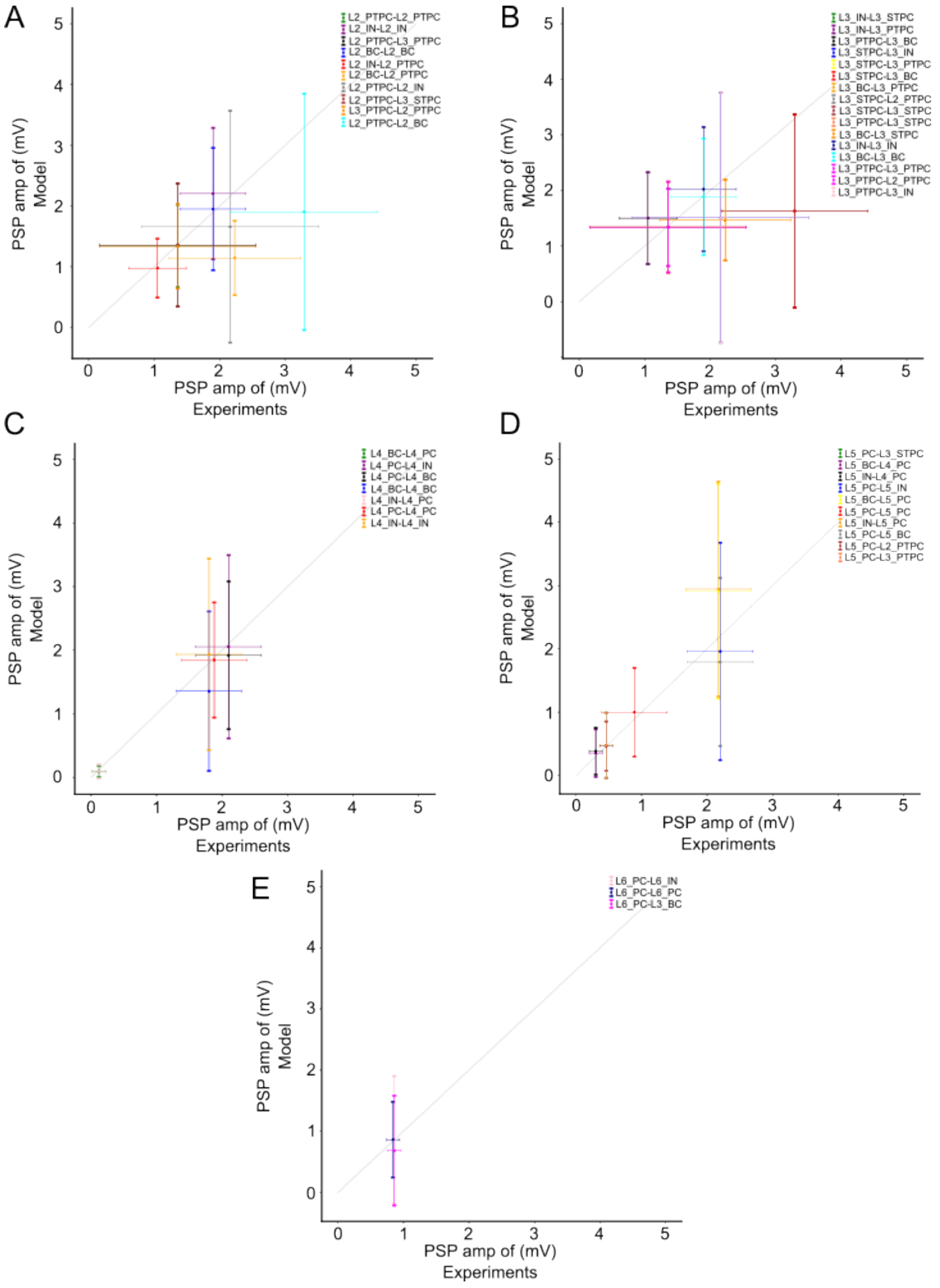


**Figure S4** Validation of synaptic parameters by calibrating gsyn to match PSP values from biological data for connections in layer 2 (A), layer 3 (B), layer 4 (C), layer 5 (D) and layer 6 (E).
